# Sex-specific effects of prenatal delta-9-tetrahydrocannabinol exposure on repetitive behavior and prefrontal cortex neuronal excitability in preadolescent rats

**DOI:** 10.64898/2026.08.10.743896

**Authors:** Sonia Aroni, Martina Di Bartolomeo, Valeria Serra, Francesco Traccis, Marco Carli, Gisella Lorrai, Marcello Serra, Paola Devoto, Pierluigi Saba, Mariangela Pucci, Roberto Frau, Claudio D’Addario, Miriam Melis

## Abstract

Cannabis is the most common illicit drug abused worldwide, and its consumption has substantially increased among pregnant women. We previously demonstrated that male preadolescent offspring prenatally exposed to Δ^9^-tetrahydrocannabinol (THC), a model of prenatal cannabinoid exposure (PCE), exhibit a mesolimbic dopamine (DA) neuron dysfunction contributing to at-risk psychotic-like (endo)phenotypes that are unmasked by acute THC exposure at preadolescence. Dysregulation of mesocortical DA signaling along with prefrontal cortex (PFC) function is also a central feature of psychotic disorders. Furthermore, studies investigating the impact of PCE on PFC in the offspring at preadolescence, a window of heightened plasticity and vulnerability, are limited. To fill this gap, we applied a multiscale analysis of mesocortical DA transmission and PFC function in PCE preadolescent offspring by integrating behavioral, neurochemical, electrophysiological, and molecular approaches. PCE enhanced spontaneous repetitive behaviors in a male-specific manner. PCE also abolished sex differences in the intrinsic excitability of PFC pyramidal neurons and Netrin-1 expression. In addition, PCE altered the expression of genes associated with endocannabinoid signaling without changing basal and THC-induced extracellular levels of DA in the PFC. Collectively, these findings demonstrate that prenatal THC exposure disrupts both proper maturation and sexual differentiation of PFC circuitry, thus extending the impact of PCE from previously described mesolimbic abnormalities to mesocortical pathway. Finally, our data identify early cortical molecular and cellular alterations that may contribute to neuropsychiatric vulnerability later in life.

**Highlights:** • Preadolescent male rats exposed *in utero* to THC display repetitive behavior

• Prenatal cannabinoid exposure (PCE) does not alter dopamine transmission in the PFC

• PCE potentiates AMPA-mediated transmission in male pyramidal cells

• PCE abolishes sex differences in Netrin-1 expression levels in the PFC

## 1. Introduction

Cannabis is the most common illicit drug abused worldwide (United Nations Office on Drugs and Crime 2024), and, probably due to its rising legalization, its misuse has substantially increased among pregnant women, who consume it to relieve anxiety, nausea, and pain (Skelton et al. 2024; Ko 2020; Zoorob and Quinlan 2024). Of note, the concentration of Δ^9^-tetrahydrocannabinol (THC), the main psychoactive ingredient of cannabis, has increased over the past 40 years, along with its associated harmfulness (Freeman et al. 2021). Importantly, THC crosses the placenta and disrupts endocannabinoid signaling (The American College of Obstetricians and Gynecologists 2024), thus contributing to deleterious effects on neurodevelopment (Metz et al. 2023; Harhangi et al. 2026).

In humans, cannabis use during pregnancy leads to adverse outcomes in the progeny (Thompson et al. 2019; Paul et al. 2021; Kleinhans et al. 2024; Munn et al. 2025; Bailey et al. 2025; Corsi et al. 2025; Azubuike et al. 2025). School-aged children exposed *in utero* to cannabis display impairments in problem-solving, memory, and attention, as well as increased anxiety and depressive symptoms (Thompson et al. 2019; Shukla and Doshi 2025). Enhanced impulsivity and increased risk of developing attention deficit hyperactivity disorder (ADHD) and autism spectrum disorders (ASD) are also associated with maternal cannabis use (Wu et al. 2011; Pinky et al. 2023; Nashed et al. 2021; Tadesse et al. 2024).

In rodents, prenatal cannabinoid exposure (PCE) alters emotional processing in offspring by disrupting the development and function of the mesocorticolimbic dopamine (DA) system, a pivotal neural circuit for modulating reward, motivation, learning, and emotional processing (Navarro et al. 2024; Jenkins et al. 2025). In particular, DA neurons arising from the ventral tegmental area (VTA) and projecting to the nucleus accumbens (NAc), amygdala, hippocampus, and prefrontal cortex (PFC) are crucial for encoding and modulating the abovementioned functions (Elum et al. 2024; Solié et al. 2022; Hou et al. 2024), which are affected by PCE (Frau et al. 2019b). PCE leads to an impairment of the emotional reactivity of juvenile and adult male offspring (Weimar et al. 2020; Sarikahya et al. 2022, 2023; Vargish et al. 2017; Bara et al. 2018; Frau et al. 2019b; Traccis et al. 2021; Sagheddu et al. 2021; Serra et al. 2024; Ellis et al. 2022) as well as an increase of repetitive behaviors in socially dominant mice (Mari et al. 2025). Furthermore, PCE male offspring exhibit increased excitability of pyramidal neurons in the PFC at adolescence and adulthood (Bara et al. 2018; Di Bartolomeo et al. 2025), accompanied by persistent impaired social behavior (Bara et al. 2018; Weimar et al. 2020; Pham et al. 2025) and a more pronounced voluntary opioid drug-seeking in adulthood compared to female littermates (Spano et al. 2007; Luján et al. 2024). In parallel, PCE induces sex-dependent alterations in glutamatergic, GABAergic, and dopaminergic signaling in the PFC at adulthood, suggesting a dysregulation of the excitatory/inhibitory (E/I) balance (Sarikahya et al. 2023). PCE also elicits a male-specific hyperdopaminergic state and deficits in pre-pulse inhibition (PPI) of the startle reflex following an acute stressor or a single exposure to THC in preadolescent offspring (Frau et al. 2019b; Sagheddu et al. 2021; Traccis et al. 2021; Serra et al. 2024). These PCE-driven and sex-specific changes may account for repetitive behaviors and stereotypies observed in both ASD patients and animal models of ASD (Fuccillo 2016; DiCarlo and Wallace 2022). PCE neurodevelopmental effects appear to display sexual specificity (Loo et al. 2024): while male preadolescents manifest pronounced behavioral anomalies along with mesocorticolimbic hyperexcitability, their female counterparts (Ellis et al. 2022) appear resilient (Traccis et al. 2021; Bara et al. 2018; Mari et al. 2025; Frau et al. 2019b; Sandini et al. 2023).

Despite this evidence, it remains unclear whether PCE-induced alterations in DA and endocannabinoid system (ECS) components within the PFC are already detectable during preadolescence, a developmental period that precedes the full emergence of adolescent and adult behavioral abnormalities, and that represents a critical window for prefrontal circuit maturation. This question is particularly relevant in light of the marked sex-specific effects of PCE, with males often showing vulnerability. Whether early PFC alterations emerge in a sex-specific manner and are associated with repetitive behaviors, a behavioral domain linked to prefrontal-striatal and broader cortico-striatal circuit dysfunction, remains poorly understood (Fuccillo 2016; DiCarlo and Wallace 2022). To address this gap, we investigated whether PCE induces sex-related cellular, molecular, and transcriptional changes in DA- and ECS- related components within the preadolescent PFC, and whether these changes are associated with the emergence of repetitive behaviors in the offspring.

## 2. Materials and Methods

### 2.1 Subjects and treatments

#### 2.1.1 Drugs

Δ^9^-tetrahydrocannabinol (THC) resin was purchased from THC PHARM GmbH (Frankfurt, Germany), dissolved in ethanol at 20% final concentration, and then sonicated for 30 minutes. THC was emulsified in 1-2% Tween®80 and then dissolved in sterile saline (0.9% NaCl).

#### 2.1.2 Animals and treatments

Primiparous female Sprague Dawley rats (Envigo) were used as mothers and single housed during pregnancy. THC or vehicle was administered (2 mg/kg/mL, subcutaneously (s.c.), once daily) from gestational day (GD) 5 to GD 20, as previously described (Frau et al. 2019b). This dose was selected because repeated administration does not induce overt behavioral responses or cannabinoid tolerance (Wiley et al. 2007), nor does it substantially affect maternal or non-maternal behavior, or offspring body weight (Frau et al. 2019b). Furthermore, it is equivalent in rodent plasma concentrations (8.6-12.4 ng/mL) to human recreational cannabis smokers (7% THC) 0-22 hours post-inhalation (13-63 ng/mL) (Schwope et al. 2011; Klein et al. 2011). The offspring were weaned at postnatal day (PND) 21 and were housed in a climate-controlled animal room (21±1°C; 60% humidity) under a normal 12-hour light-dark cycle (lights on at 7:00 a.m.) with water and food available *ad libitum* until the experimental day (PND 24-29). For the experiments we used male and female offspring, and to control for litter effects, we did not use more than two offspring from each litter for the same experiment. All procedures were performed in accordance with the European legislation (EU Directive, 2010/63) and were approved by the Animal Ethics Committees of the University of Cagliari and by Italian Ministry of Health (auth. n. 636/2022-PR). All possible efforts were made to minimize animal pain and discomfort and to reduce the number of experimental subjects.

### 2.2 Marble burying test

The marble burying test was performed in transparent rat cages (54×34.5×20 cm) under a dim light (30 lux) (Bratzu et al. 2023). Male and female rats (PND 24-29) were individually placed into the cage containing 5 cm of litter bedding and 15 glass marbles on the box floor arranged in 5 rows of 3 marbles each. At the beginning of the test session, each animal was placed in a marble-free area of the experimental cage and allowed to freely explore the cage for 30 min. While a video camera located on the room ceiling monitored the animal’s activity. The animals were video-recorded during the test and then scored offline by blinded observers. Behavioral measures included the number of marbles buried at least two-thirds of their depth (Angoa-Pérez et al. 2013), the duration of the digging and the grooming.

### 2.3 Cerebral microdialysis

As previously described (Traccis et al. 2021), for cerebral microdialysis experiments, male and female rats were anesthetized with Equithesin (0.97 g pentobarbital, 2.1 g MgSO_4_, 4.25 g chloral hydrate, 42.8 ml propylene glycol, 11.5 ml 90% ethanol in 100 ml; 5 ml/kg, intraperitoneally (i.p.)) and stereotaxically implanted with home-made vertical microdialysis probes (AN 69-HF membrane, Hospal-Dasco; cut-off 40 kDalton, 4 mm dialyzing membrane length) in the medial prefrontal cortex (mPFC; from bregma: anterior-posterior, +2.5; lateral, 0.5; ventral, -6; (Paxinos and Watson 2013)). The day after probe implantation, artificial cerebrospinal fluid solution (aCSF; 147 mM NaCl, 4 mM KCl, 1.5 mM CaCl_2_, 1 mM MgCl_2_, pH 6–6.5) was perfused through the microdialysis probes at a constant rate of 1.1 μL/min via a CMA/100 microinjector pump (Carnegie Medicine, Stockholm, Sweden) in freely moving animals. Samples were collected every 20 minutes and analyzed for dopamine (DA) and 3,4-dihydroxyphenylacetic acid (DOPAC) content by high-performance liquid chromatography (HPLC) coupled with electrochemical detection, as previously described (Devoto et al. 2003). When a stable baseline was obtained (three consecutive samples with a variance not exceeding 15%), THC (2.5 mg/kg/2 mL) was administered (i.p.), and sample collection continued for 2 hours. On completion of the experiments, the rats were euthanized with an Equithesin overdose, and the brains were removed and sectioned using a cryostat (Leica CM3050 S) into 40 μm thick coronal slices to verify the anatomical locations of microdialysis probes.

### 2.4 *Ex vivo* electrophysiological recordings

We prepared coronal prefrontal cortex (PFC) slices (300 μm) from PND 24-29 male and female offspring. Rats were anaesthetized with isoflurane until loss of righting reflex, then the brain was rapidly removed, and PFC slices were obtained using an ice-cold sucrose-based solution saturated with 95% O_2_/5% CO_2_, containing in mmol/L: 87 NaCl, 75 sucrose, 25 glucose, 5 KCl, 21 MgCl_2_, 0.5 CaCl_2_ and 1.25 NaH_2_PO_4_ (Kasanetz et al. 2013; Frau et al. 2019a) with a vibratome (Leica VT 1000S). Immediately after cutting, slices were stored for 1 h at 32°C in an artificial cerebrospinal fluid (aCSF) at 304–306 mOsm, and contained, in mmol/L:130 NaCl, 11 glucose, 2.5 KCl, 1.2 MgCl_2_, 2.4 CaCl_2_, 23 NaHCO_3_, 1.2 NaH_2_PO_4_, and were equilibrated with 95% O_2_/5% CO_2_. Slices were then stored in aCSF at room temperature until recording. Cells were visualized with an upright microscope with infrared illumination (Axioskop FS 2 plus; Zeiss), and whole-cell patch-clamp recordings were made by using an Axopatch 200 B amplifier (Molecular Devices). Pyramidal neurons in the prelimbic portion of the PFC, were identified by their pyramidal shape, the presence of a prominent apical dendrite, and the distance from the pial surface (layers V/VI). The stimulating electrode was placed in layers II/III of the prelimbic cortex. Current-clamp and voltage-clamp recordings of evoked inhibitory postsynaptic currents (IPSCs) were made with electrodes (resistance of 4–6 MΩ) filled with a solution containing the following (in mM): 144 KCl, 10 HEPES buffer, 3.45 BAPTA, 1 CaCl_2_, 2.5 Mg_2_ATP and 0.25 Mg_2_GTP, pH 7.3–7.4, 283–285 mOsm. Current-clamp experiments were performed in the absence of any pharmacological blocker (regular aCSF), the membrane potential was held near -65 mV, and evoked firing was measured using depolarizing current steps (0.4 s) from 0 to 400 pA. For voltage-clamp recordings, the membrane potential was held at -80 mV, and all GABA_A_ IPSCs were recorded in the presence of 6-cyano-2,3-dihydroxy- 7-nitro-quinoxaline (10 μM) and AP-5 (100 μM) to block AMPA- and NMDA-mediated synaptic currents. Voltage-clamp recordings of evoked excitatory postsynaptic currents (EPSCs) were made with electrodes (resistance of 4–6 MΩ) filled with a solution containing the following (in mM): 117 Cs methanesulfonic acid, 20 HEPES buffer, 0.4 EGTA, 2.8 NaCl, 5 TEA-Cl, 0.1 spermine, 2.5 Mg_2_ATP and 0.25 Mg_2_GTP, pH 7.2–7.4, 282–284 mOsm. Picrotoxin (100 μm) was added to the aCSF for recording, to block GABA_A_ receptor-mediated IPSCs. In addition, random experiments were performed with an internal solution that contained biocytin (0.2%) to allow for subsequent immunocytochemical detection of Netrin-1. Experiments were begun only after series resistance had stabilized (typically 15–40 MΩ). Data were filtered at 2 kHz, digitized at 10 kHz and collected online with acquisition software (pClamp 10.2, Molecular Devices).

### 2.5 Immunofluorescence analysis

After electrophysiological recordings, 240 µm-thick coronal sections of the medial prefrontal cortex (mPFC) were post-fixed in 4% paraformaldehyde (PFA) in 0.2 M phosphate buffer (PB) for 24 hours, rinsed three times in 1X PBS, and stored in 0.05% sodium azide in 1X PBS at 4°C until use. To detect biocytin and Netrin-1 immunostaining, antigen retrieval was performed by incubating sections in sodium citrate buffer (pH 6) at 95°C for 25 minutes. The free-floating sections were then rinsed in 0.1 M PB and blocked overnight at 4°C in a solution containing 10% normal goat serum (Vector, UK), 0.05% bovine serum albumin, and 0.5% Triton X-100 in 0.1 M PB. Subsequently, the sections were incubated at 4°C with a rabbit polyclonal anti-Netrin-1 primary antibody (1:600, Alomone Labs, #ANR-121, 120 h), rinsed three times in 0.1 M PB, and then incubated at room temperature for 3 hours with AlexaFluor-488 goat anti-rabbit IgG secondary antibodies (1:200, Jackson ImmunoResearch, UK, #111-545-003) and streptavidin-594 (1:200, Jackson ImmunoResearch, UK, #016-580-084). Thereafter, the sections were incubated for 10 minutes in 4′,6-diamidino-2-phenylindole (DAPI, 1:10,000, Merck, Italy, D9542) for nuclear visualization, rinsed in 0.1 M PB, and mounted onto Superfrost glass slides using Mowiol® mounting medium. Negative controls, in which primary or secondary antibodies were omitted, produced no labelling (data not shown). Single-wavelength images (14-bit depth) were acquired using a ZEISS Axio Scan Z1 slide scanner (Zeiss, Germany). Brain sections were captured at 40x magnification to encompass the entire mPFC. ImageJ software (National Institutes of Health, USA) was used to visualize biocytin-filled cells and assess their spatial overlap with Netrin-1 immunoreactivity.

### 2.6 Molecular biology analysis

#### 2.6.1 Gene expression analysis by Quantitative Real Time-Polymerase Chain Reaction (qRT-PCR)

Total RNA was extracted from dissected rat PFC tissue using Qiazol® Reagent (Qiagen, Hilden, DE) following the manufacturer’s protocol. Concentration and purity were assessed using a NanoDrop 2000c UV-Vis Spectrophotometer (Thermo Fisher Scientific, Waltham, MA, United States). RNA integrity was then verified by 1% agarose gel electrophoresis. 1 µg of total RNA was converted into cDNA by SensiFAST^TM^ cDNA Synthesis kit (Bioline Reagents, London, UK). Real-time quantitative polymerase chain reaction (RT-qPCR) was performed using the SensiFAST™ SYBR^®^Lo-ROX Kit (Bioline Reagents, London, UK) on a DNA Engine Opticon 2 Continuous Fluorescence Detection System (MJ Research). For accurate target quantification, amplification curves of each PCR reaction were analyzed to determine the threshold cycle (Ct) value, representing the point at which fluorescence exceeded background levels during the early exponential phase, as previously described (Di Bartolomeo et al. 2021, 2023, 2025). Relative target quantities were calculated based on Ct value differences between samples. Post-amplification melting curve analysis (60-95°C) was performed to confirm product specificity (Lyon 2001). Relative mRNA expression levels were calculated using the delta-delta Ct (ΔΔCt) method and converted to 2^−ΔΔCt^ for statistical analysis (Livak and Schmittgen 2001). All data were normalized to two endogenous reference genes: glyceraldehyde-3-phosphate dehydrogenase (GAPDH) and 18S ribosomal RNA (18S). Sequences of the primers used for the amplification are reported in **Table 1**.

**Table 1.**
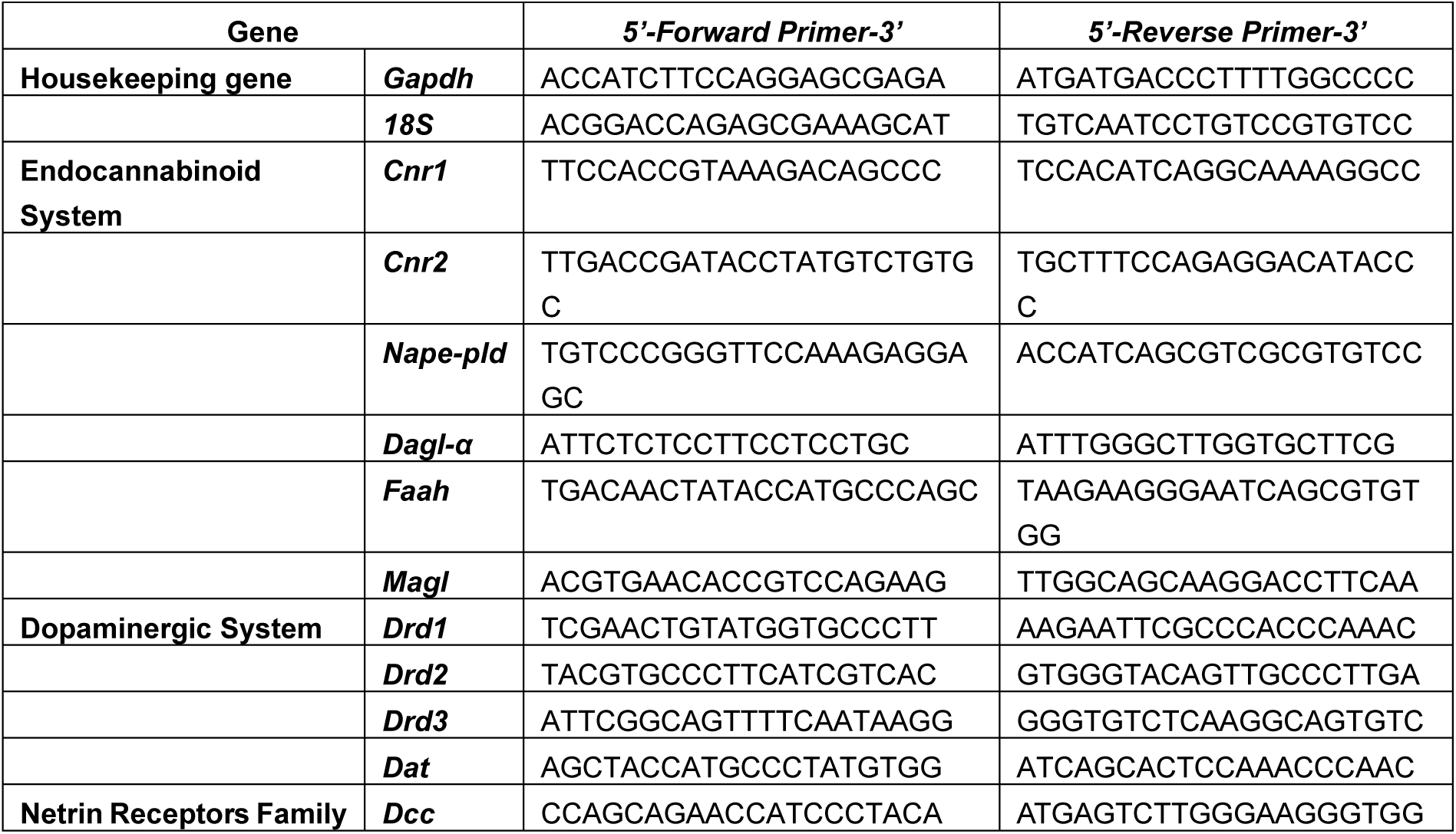
List of primers used for quantitative real-time polymerase chain reaction (qRT-PCR).

#### 2.6.2 DNA methylation analysis by pyrosequencing

To analyze methylation levels of *Magl* (encoding for the Monoacylglycerol lipase) and *Dat* (encoding for the DA transporter), genomic DNA was extracted from rat PFC using Qiazol® Reagent (Qiagen, Hilden, Germany). Concentration and purity of each DNA sample were detected by NanoDrop 2000c UV-Vis Spectrophotometer (Thermo Fisher Scientific, Waltham, MA, United States). Each purified DNA sample was subjected to bisulfite modification using the EZ DNA Methylation-Gold^TM^ Kit (Zymo Research, Orange, CA, US), according to the manufacturer’s protocol. Pyrosequencing was used to quantify the methylation levels of individual CpG sites of *Magl* and *Dat* gene regulatory regions. Bisulfite-converted DNA was amplified using the PyroMark PCR Kit (Qiagen, Hilden, Germany) with biotin-labeled primers following the manufacturer’s protocol (Di Bartolomeo et al. 2021, 2023, 2024, 2025). The PCR program consisted of an initial denaturation step at 95°C for 15 minutes, followed by 45 cycles of denaturation (94°C, 30 s), annealing (56°C, 30 s), and extension (72°C, 30 s), with a final extension at 72°C for 10 minutes. PCR product specificity was confirmed by gel electrophoresis. Pyrosequencing was conducted on a PyroMark Q48 Autoprep system using PyroMark Q48 Advanced Reagents (Qiagen, Hilden, Germany) according to the manufacturer’s instructions. A specific PyroMark CpG assay (Qiagen, Hilden, Germany), targeting 7 CpG sites, was used to analyze the rat *Magl* regulatory region, while primers for the rat *Dat* gene, targeting 6 CpG sites, were designed using the PyroMark Assay Design software v2.0 (Qiagen, Hilden, Germany). Methylation quantification was performed using the PyroMark Q48 Autoprep 2.4.2 software, calculating methylation percentages via the mC/(mC+C) ratio (mC=methylated cytosine, C=unmethylated cytosine) for each CpG site to enable quantitative comparisons. Data were expressed as individual CpG site methylation percentages and as the mean methylation of all analyzed sites.Sequence and primer information are shown in **Table 2**.

**Table 2.**
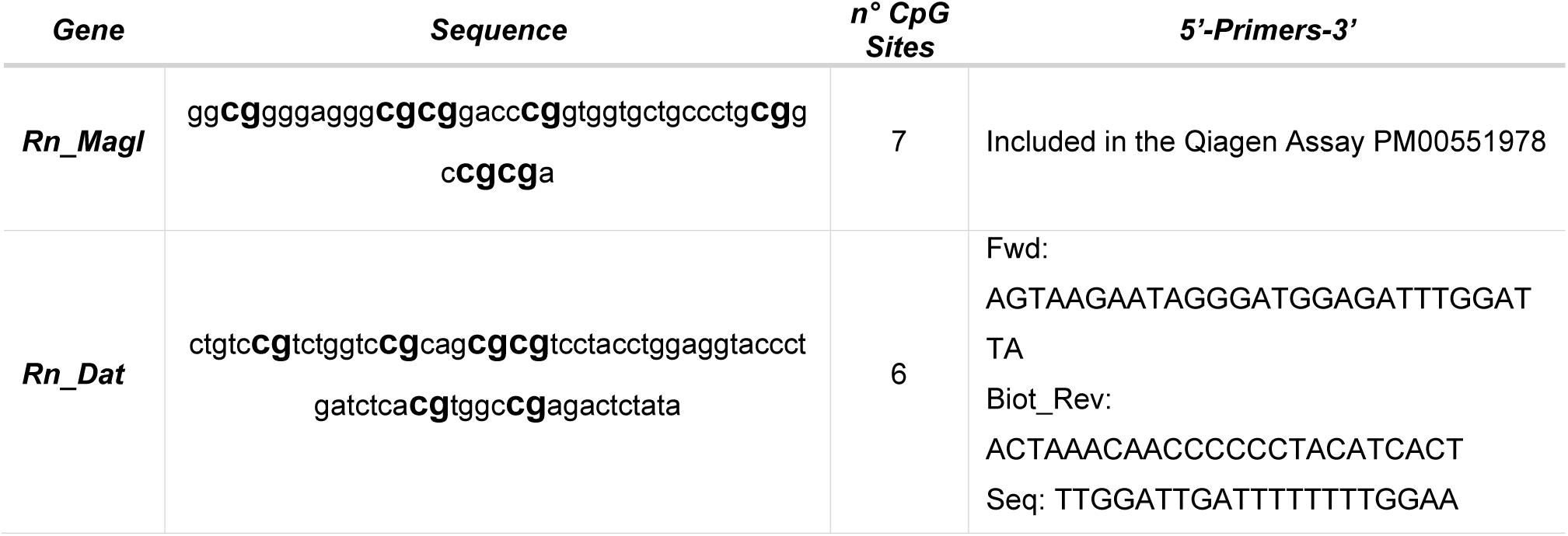
Details of sequences and primers employed for DNA methylation analysis by pyrosequencing. Rn = Rat; CpG = C-phosphate-G; Fwd = forward primer; Biot_Rev = biotinylated reverse primer; Seq = sequencing primer. Bold text indicates CpG sites analyzed.

### 2.7 Netrin-1 and tyrosine hydroxylase immunohistochemistry

#### 2.7.1 Tissue preparation

At PND 25-30, rats were deeply anesthetized with isoflurane and transcardially perfused with saline, followed by 4% paraformaldehyde in 0.1 M PB. Brains were then removed and post-fixed overnight at 4°C in the same solution. The following day, brains were rinsed three times in 1X PBS and then cryoprotected in a 30% sucrose solution in 1X PBS at 4°C until sinking (approximately 48 hours). Subsequently, the tissue was rapidly frozen by immersion in isopentane cooled in dry ice. Afterward, brains were embedded in optimal cutting temperature (OCT) compound and coronally cut on a cryostat (LEICA CM3050S, Leica Biosystems) to yield sections (thickness, 40 µm) suited for immunohistochemistry (IHC) processing. For each rat, at least two coronal sections representative of the mPFC and nucleus accumbens (NAc), containing both the core and shell substructures were collected based on stereotaxic coordinates ranging from +1.50 mm to +2.50 mm relative to bregma. These coordinates were referenced from the Atlas of the Postnatal Rat Brain in Stereotaxic Coordinates (Khazipov et al. 2015).

#### 2.7.2 Reaction protocol, image acquisition, and density analysis

For sections used for Netrin-1 IHC, antigen retrieval was performed by placing sections in a sodium dodecyl sulfate (SDS) 1% solution for 5 minutes, a protocol optimized by Salameh et al., (Salameh et al. 2018). Free-floating sections were rinsed three times in 0.1 M PB and blocked in a solution containing 10% normal goat serum (Cell Signaling Technology, Danvers, MA, USA) and 0.25% Triton X-100 in 0.1 M PB at room temperature (2 h). Thereafter, sections were incubated at 4°C with either the rabbit polyclonal primary antibody anti-tyrosine hydroxylase (TH, 1:1000, Merck, Darmstadt, Germany, #AB152, 48 h) or rabbit polyclonal primary antibody anti-Netrin-1 (1:600, Alomone Labs, Jerusalem, Israel, #ANR-121, 120 h), rinsed three times in 0.1 M PB, and then incubated with the secondary antibody, Alexa Fluor^®^488 Conjugate anti-rabbit IgG (1:1000, Cell Signaling Danvers, MA, USA, #4412) in 0.1 M PB at room temperature (3 h). Afterward, sections were rinsed three times in 0.1 M PB and mounted onto super-frost glass slides using mounting medium containing DAPI (Abcam, MA, USA, ab104139) for cell nuclei visualization. Images were obtained with a ZEISS Axio Scan Z1 slide scanner (Zeiss, Oberkochen, Germany). Brain sections were captured at 20X magnification (Objective: Plan-Apochromat 20x/0.8 M27) to acquire the whole mPFC and NAc from both hemispheres. The ImageJ software (v.1.54, National Institutes of Health, Bethesda, Maryland, USA) was used to measure the density of Netrin-1 and TH signals in both brain areas. Images were converted to 16-bit, background-subtracted, and the signal density was quantified in regions of interest representative of each brain area, with dimensions of 150 x 150 µm. No significant differences in the relative density analysis were found between NAc subregions, therefore, measurements from core and shell immunoreactivity levels were averaged. Analyses were conducted blinded to the treatment of each animal. No significant differences in the relative density of TH and Netrin-1 immunoreactivity were found between the two hemispheres of the two sections, therefore values from different ventro-lateral and antero-posterior levels were averaged.

### 2.8 Statistical analysis

All data were presented as box-and-whisker plots showing single values (min to max) or as mean ± standard error of the mean (S.E.M.). Statistical differences between the experimental groups were evaluated using GraphPad Prism® 9 (Graph-Pad Software, San Diego, CA). The nonparametric Mann–Whitney test (M-W test) was used for analyzing data from behavior, microdialysis, *ex vivo* electrophysiology, and gene expression experiments. 2-way ANOVA for repeated measure (RM) followed by Sidak’s *post-hoc* test was used when appropriated for microdialysis and *ex vivo* electrophysiology data. 2-way ANOVA followed by Sidak’s *post-hoc* test was used for IHC data. DNA methylation at each CpG site as well as in the average of the CpG sites under study was analyzed using the M-W test and Holm–Šídák correction was used for the multiple comparisons as recommended. Statistical outliers were identified with Grubb’s test (α = 0.05) and excluded from the analysis. p-values < 0.05 were considered to be statistically significant.

## 3. Results

### 3.1 PCE enhances repetitive behavior in male preadolescent rats

Prenatal cannabinoid exposure (PCE; **Fig. 1a**) enhanced the number of marbles buried by male offspring in the marble burying test (U=34, p=0.0320, M-W test; **Fig. 1b**), as well as the overall time spent digging (U=19.50, p=0.0033, M-W test; **Fig. 1c**), without affecting grooming (U=63.50, p=0.9393, M-W test; **Fig. 1d**), indicating an enhanced repetitive/perseverative behavior. In contrast, PCE did not affect marble burying behavior in females (marbles buried: U=66, p=0.1931, M-W test; **Fig. 1e**; digging: U=55, p=0.0747, M-W test; **Fig. 1f**; grooming: U=88, p=0.7807, M-W test; **Fig. 1g**), thus supporting a sex-specific effect of prenatal THC exposure.

**Figure 1.**
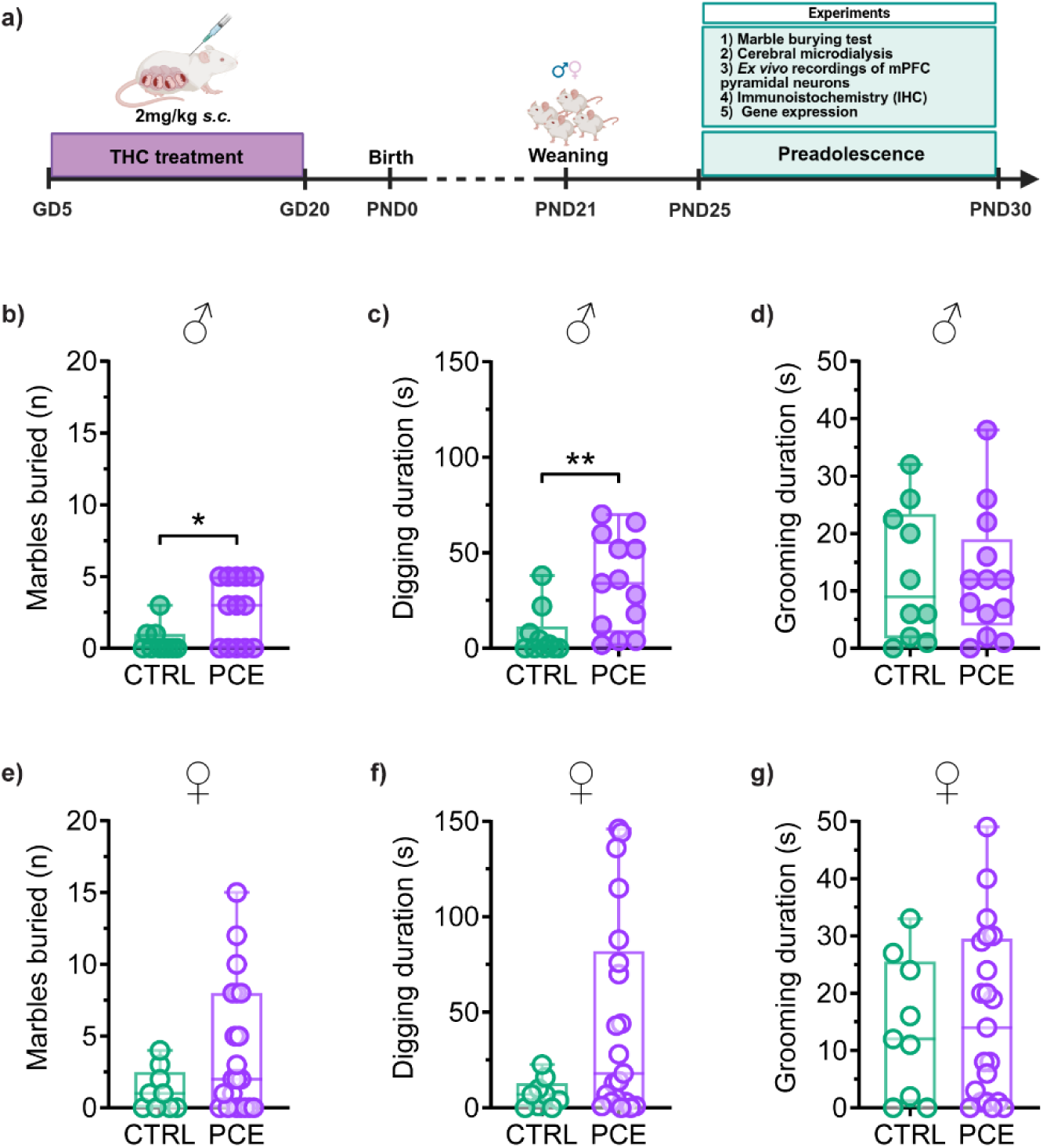
PCE only enhances repetitive behavior in male preadolescent rats. (**a**) Timeline of PCE treatment and experiments. GD=gestational day; PND=postnatal day. Figure created with BioRender.com. (**b-d**) In males, PCE increased the total number of marbles buried (**b**) and the overall time spent digging (**c**), without affecting grooming (**d**; n=10-13/group). (**e-g**) PCE female progeny displayed no changes in the number of marbles buried (**e**), digging (**f**) and grooming (**g**; n=9-21/group). Data are expressed as box-and-whisker plots with single values (min to max). **p<0.01; *p<0.05.

### 3.2 PCE does not affect mesocortical dopamine transmission in preadolescent offspring

Given that dopamine (DA) transmission within the prefrontal cortex (PFC) acts as a critical tuner for repetitive behaviors (Denys et al. 2004; Yin et al. 2024), we carried out *in vivo* cerebral microdialysis experiments in the medial PFC (mPFC) to reveal potential PCE effects (**Fig. 2**). PCE did not change either basal (U=29, p=0.3401, M-W test; **Fig. 2a**) or THC-induced increase in extracellular DA in males (p=0.9650, 2-way RM ANOVA, interaction: F_(8,144)_=0.2997; **Fig. 2b**). Likewise, extracellular levels of 3,4-dihydroxyphenylacetic acid (DOPAC), an indirect index reflecting DA synthesis and turnover (Goldstein et al. 2018), displayed a similar profile (basal: U=39, p=0.2816, M-W test; **Fig. 2c**; THC-induced: p=0.8357, 2-way RM ANOVA, interaction: F_(8,171)_=0.5261; **Fig. 2d**). Similarly, female offspring displayed no differences between the groups when basal DA (U=25, p=0.3089, M-W test; **Fig. 2e**) or THC-induced alterations were evaluated (p=0.7448, 2-way RM ANOVA, interaction: F_(8,132)_=0.6378; **Fig. 2f**). Accordingly, no changes were found in female DOPAC profile (basal: U=26, p=0.2135, M-W test; **Fig. 2g**; THC challenge: p=0.9820, 2-way RM ANOVA, interaction: F_(8,144)_=0.2426; **Fig. 2h**).

**Figure 2.**
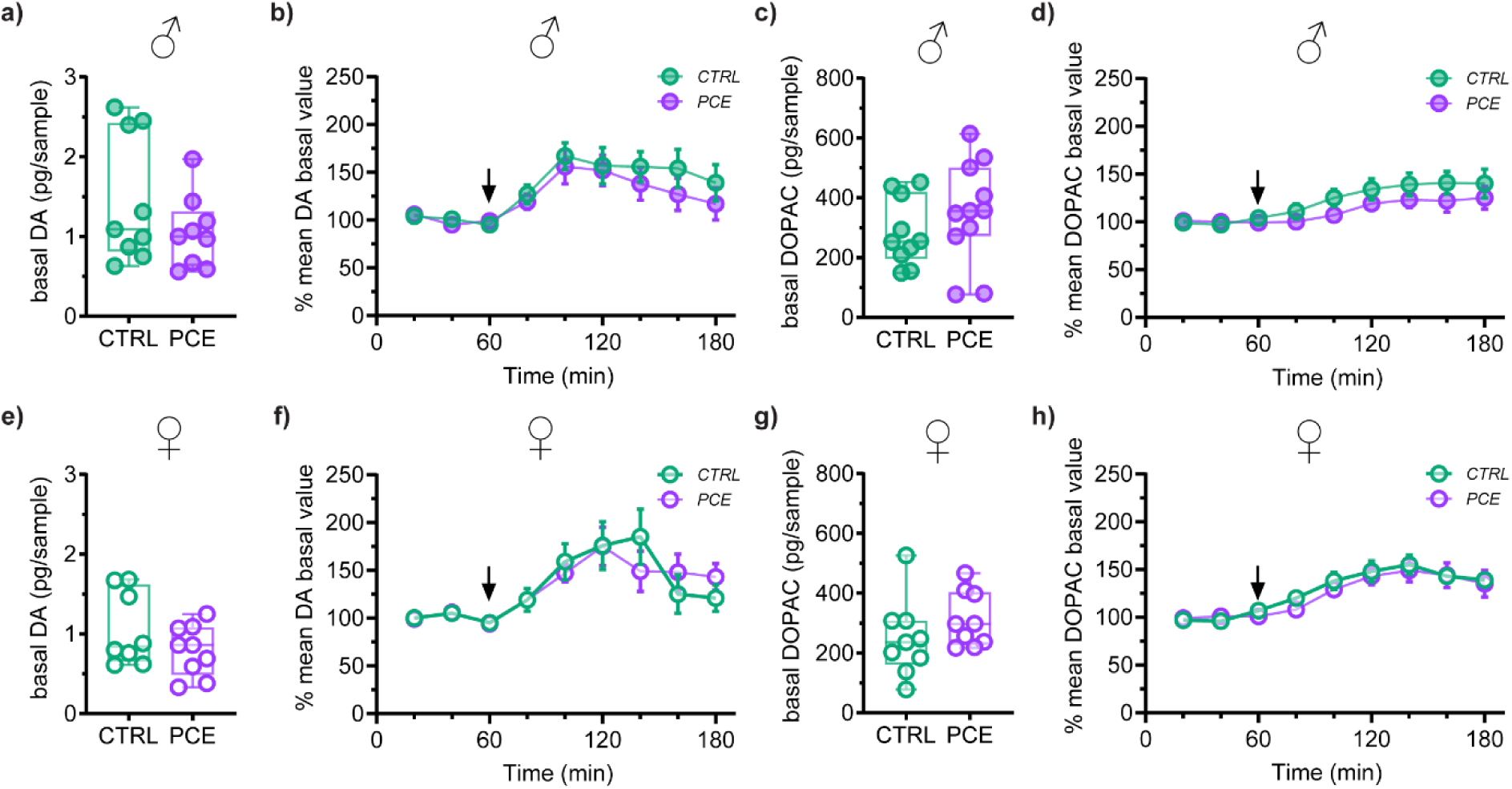
PCE does not affect dopamine transmission in the PFC. (**a**) Basal dopamine (DA) levels (pg) measured in the mPFC of male rats (n=9/group). (**b**) Time course of the effect of acute THC (2.5 mg/kg/2 mL) on extracellular DA concentrations in the mPFC of male offspring (n=9/group). (**c**) Basal 3,4- dihydroxyphenylacetic acid (DOPAC) levels (pg) measured in the mPFC of male rats (n=10-11/group). (**d**) Time course of the effect of acute THC (2.5 mg/kg/2 mL) on extracellular DOPAC levels in the mPFC of male offspring (n=10-11/group). (**e**) Basal DA levels (pg) measured in the mPFC of female animals (n=8-9/group). (**f**) Time course of the effect of acute THC (2.5 mg/kg/2 mL) on extracellular DA concentrations in the mPFC of female progeny (n=8-9/group). (**g**) Basal DOPAC levels (pg) measured in the mPFC of female animals (n=9/group). (**h**) Time course of the effect of acute THC (2.5 mg/kg/2 mL) on extracellular DOPAC levels in mPFC of female progeny (n=9/group). Basal DA and DOPAC level data are expressed as box-and-whisker plots with single values (min to max). The arrow indicates the time of acute THC administration. DA and DOPAC time course data are represented as mean ± S.E.M.

### 3.3 PCE enhances PFC pyramidal neuron excitability in a male-specific manner at pre-adolescence

PCE enhances male PFC pyramidal neuron excitability from adolescence (Di Bartolomeo et al. 2025) throughout adulthood (Bara et al. 2018). To assess whether this effect was already evident at preadolescence, we performed *ex vivo* electrophysiological recordings from pyramidal neurons located in layers V/VI of the prelimbic portion of the PFC (**Fig. 3a**). In males, PCE increased excitability of PFC pyramidal neurons as measured as a higher number of evoked action potentials when the same current is somatically injected (p<0.0001, 2-way RM ANOVA, interaction: F_(10,280)_=4.114; **Fig. 3b**). When we evaluated their intrinsic properties, we found no difference in latency of the first evoked action potential (U=155, p=0.1616, M-W test; **Fig. 3c**), voltage threshold (V_threshold_; U=175.5, p=0.3886, M-W test; **Fig. 3d**), and resting membrane potential (RMP; U=186, p=0.4027, M-W test; **Fig. 3e**). In females, PCE did not alter PFC pyramidal neuron excitability (p=0.9999, 2-way RM ANOVA, interaction: F_(10,285)_=0.09105; **Fig. 3f**) and their intrinsic properties: latency of the first evoked action potential (U=94, p=0.3133, M-W test; **Fig. 3g**), voltage threshold (V_threshold_; U=108, p=0.4618, M-W test; **Fig. 3h**), and resting membrane potential (RMP; U=137, p=0.8182, M-W test; **Fig. 3i**).

**Figure 3.**
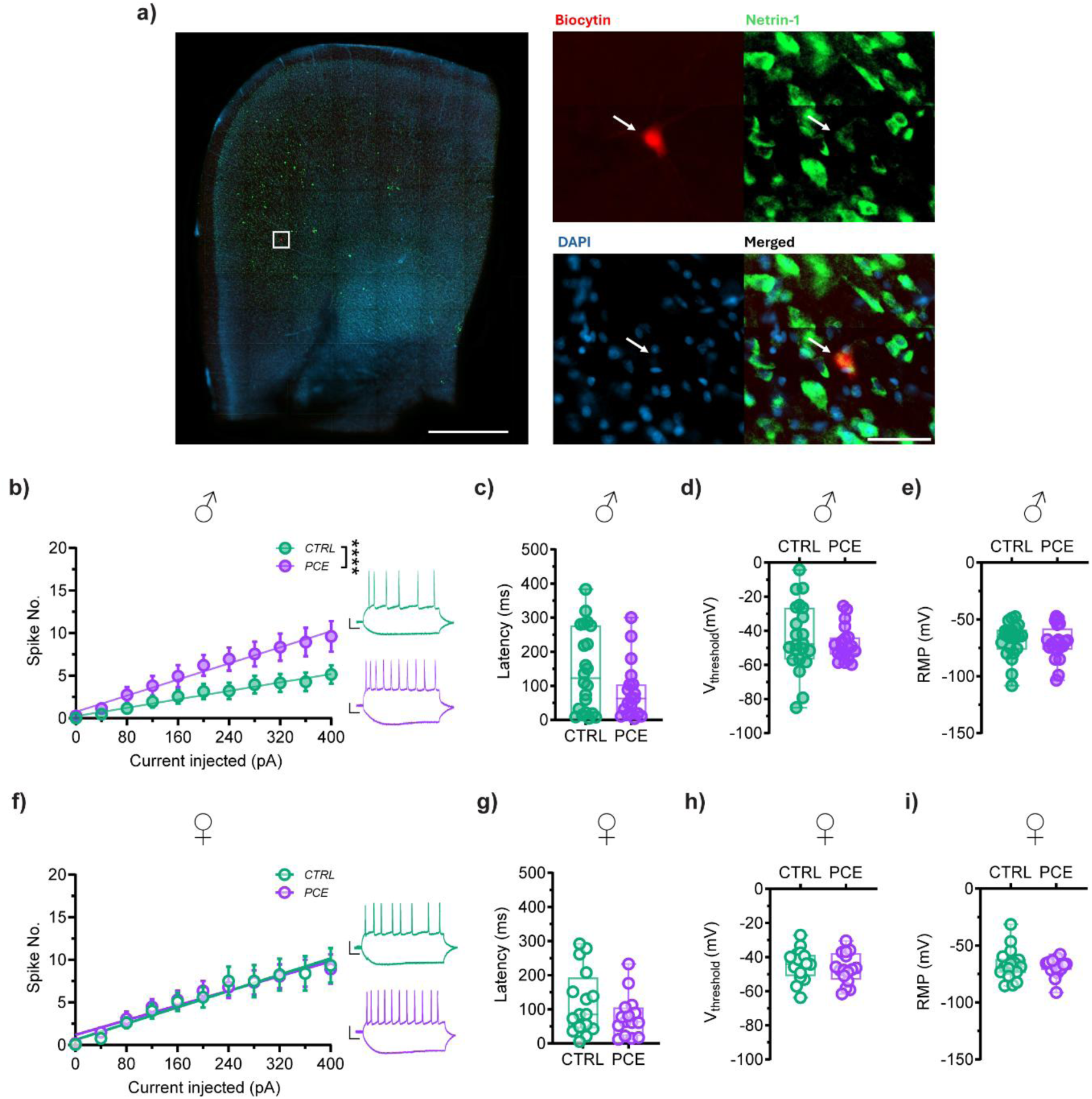
PCE enhances the excitability of male pyramidal neurons at preadolescence. (**a**) Representative fluorescent image of biocytin and Netrin-1 positive cells in a coronal brain hemisection containing the mPFC, counterstained for DAPI (left panel). White square indicates the ROI with a biocytin positive cell (left panel). Scale bar, 1000 μm. High magnifications of white ROI for labeling: biocytin (top left, red), netrin-1 (top right, green), DAPI (bottom left, blue), merged (bottom right). Scale bar, 50 μm. (**b**) Male PCE pyramidal neurons exhibited increased excitability in response to somatically injected current (interaction current x treatment ****p<0.0001; 2-way RM ANOVA) when compared to CTRL neurons (n_cells_=15/group; n_rats_=4-5/group). Insets show representative traces of evoked action potentials (APs) in response to the first and last current steps injected; calibration bar: 50 ms, 25 mV. Data are presented as mean ± S.E.M. (**c-e**) Box-and-whisker plots with single values (min to max) show intrinsic properties of male pyramidal neurons of PCE and CTRL offspring: latency for the first evoked AP (**c**); voltage threshold (V_threshold_) (**d**); resting membrane potential (RMP) (**e;** n_cells_=19-22/group, n_rats_=8-9/group). (**f**) Females showed no difference in pyramidal neurons excitability (interaction current x treatment p=0.9999; 2-way RM ANOVA; n_cells_=17-18/group; n_rats_=7-8/group) as well as in their intrinsic properties: latency for the first evoked AP (**g**); V_threshold_ (**h**); RMP (**i**: n_cells_=16-18/group, n_rats_=7-8/group).

### 3.4 PCE affects synaptic properties of medial prefrontal cortex pyramidal neurons in preadolescent rats

The excitability of pyramidal neurons might depend on the balance between excitatory and inhibitory inputs. In males, when we computed the ratio between excitatory postsynaptic currents (EPSCs) and inhibitory postsynaptic currents (IPSCs), we found no effect of PCE (E/I ratio; U=61, p=0.2465, M-W test; **Fig. 4a**). When we examined the properties of AMPA EPSCs, we found an overall PCE-induced increased amplitude of AMPA EPSCs at -80 mV (p=0.0063, 2-way RM ANOVA, interaction: F_(3,66)_=4.488; **Fig. 4b**) without changes in paired-pulse facilitation (U=49, p=0.7045, M-W test; **Fig. 4c**). PCE also augmented the AMPA/NMDA ratio (U=8.500, p=0.0019, M-W test; **Fig. 4d**) as well as AMPA EPSC amplitude at +40 mV (U=22, p=0.0381, M-W test; **Fig. 4e**). Furthermore, PCE decreased AMPA EPSC decay time in male progeny (τ; U=10, p=0.0030, M-W test; **Fig. 3f**) without affecting either NMDA amplitude (U=41, p=0.5516, M-W test; **Fig. 4g**) or NMDA decay time (τ; males: U=29, p=0.1308, M-W test; **Fig. 4h**).

**Figure 4.**
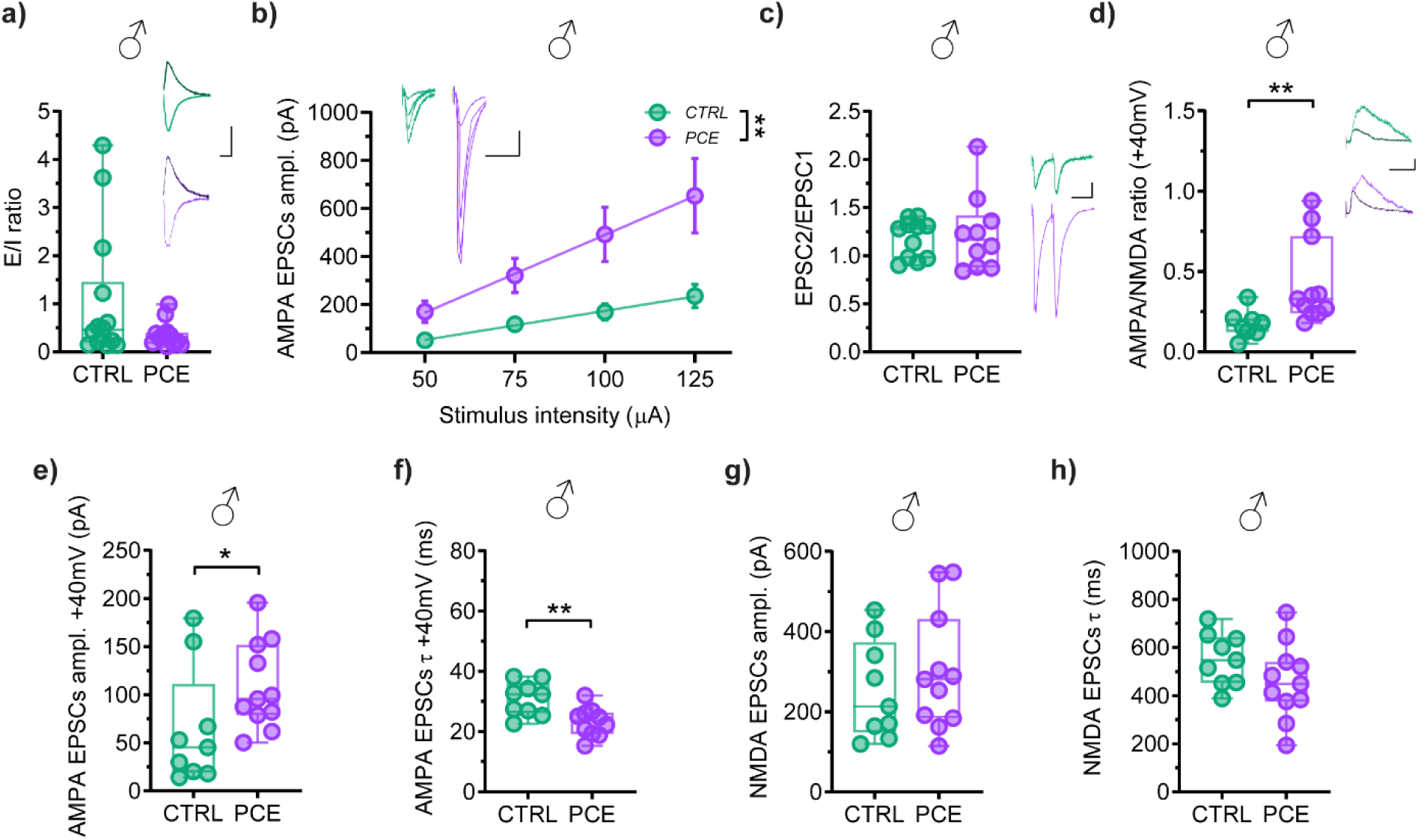
Effect of PCE on excitatory synaptic properties of PFC pyramidal neurons in preadolescent male offspring. (**a**) Plots and traces of evoked EPSCs (at -80 mV) and IPSCs (at 0 mV) recorded from PFC pyramidal neurons of CTRL and PCE male rats (p=0.2465; Mann–Whitney test; n_cells_=12-14/group, n_rats_=7/group); calibration bar: 25 ms, 200 pA. (**b**) PCE effect on input-output relationships of AMPA-mediated EPSCs in males (interaction current x treatment **p=0.0063; 2-way RM ANOVA (n_cells_=12/group; n_rats_=6-7/group). (**c**) Plots and traces of averaged paired-pulse ratio (EPSC2/EPSC1) of AMPA EPSCs of male pyramidal neurons at the highest stimulus intensity (n_cells_=10-11/group; n_rats_=6-7/group). Each symbol represents the averaged value (±S.E.M.) obtained from different cells. Insets show representative traces of AMPA EPSCs recorded from pyramidal neurons at each stimulus intensity; (**b,c**) calibration bar: 50 ms, 100 pA. (**d**) AMPA/NMDA ratio increased in PCE male offspring (**p=0.0019; Mann–Whitney test; n_cells_=8-11/group; n_rats_=4/group). Insets show representative traces of AMPA and NMDA EPSCs recorded from pyramidal neurons held at +40 mV in PFC slices; calibration bar: 25 ms, 200 pA. (**e,f**) Quantification of the data showing AMPA EPSC amplitude (**e**; *p=0.0381; Mann–Whitney test) and decay time kinetics (weighted tau, τ) (**f**; **p=0.0030; Mann–Whitney test) in CTRL and PCE male rats (n_cells_=9-11/group; n_rats_=4/group). (**g,h**) Quantification of the data showing NMDA EPSC amplitude (**g**) and τ (**h**) in CTRL and PCE male rats (n_cells_=9-11/group; n_rats_=4/group).

Female progeny did not show any change in E/I ratio (U=86, p=0.4915, M-W test; **Fig. 5a**), AMPA EPSC amplitude at -80 mV (p=0.9826, 2-way RM ANOVA, interaction: F_(3,59)_=0.05557; **Fig. 5b**) paired-pulse facilitation (U=20, p=0.0506, M-W test; **Fig. 5c**), AMPA/NMDA ratio (U=43, p=0.8868, M-W test; **Fig. 5d**), and AMPA EPSC amplitude at +40 mV (U=26, p=0.1333, M-W test; **Fig. 5e**). However, PCE female offspring also showed a decreased AMPA EPSC decay time at + 40mV (U=15, p=0.0037, M-W test; **Fig. 5f**) without effects on NMDA amplitude (U=44, p=0.7103, M-W test; **Fig. 5g**) and NMDA decay time (τ; U=31, p=0.3100, M-W test; **Fig. 5h**).

**Figure 5.**
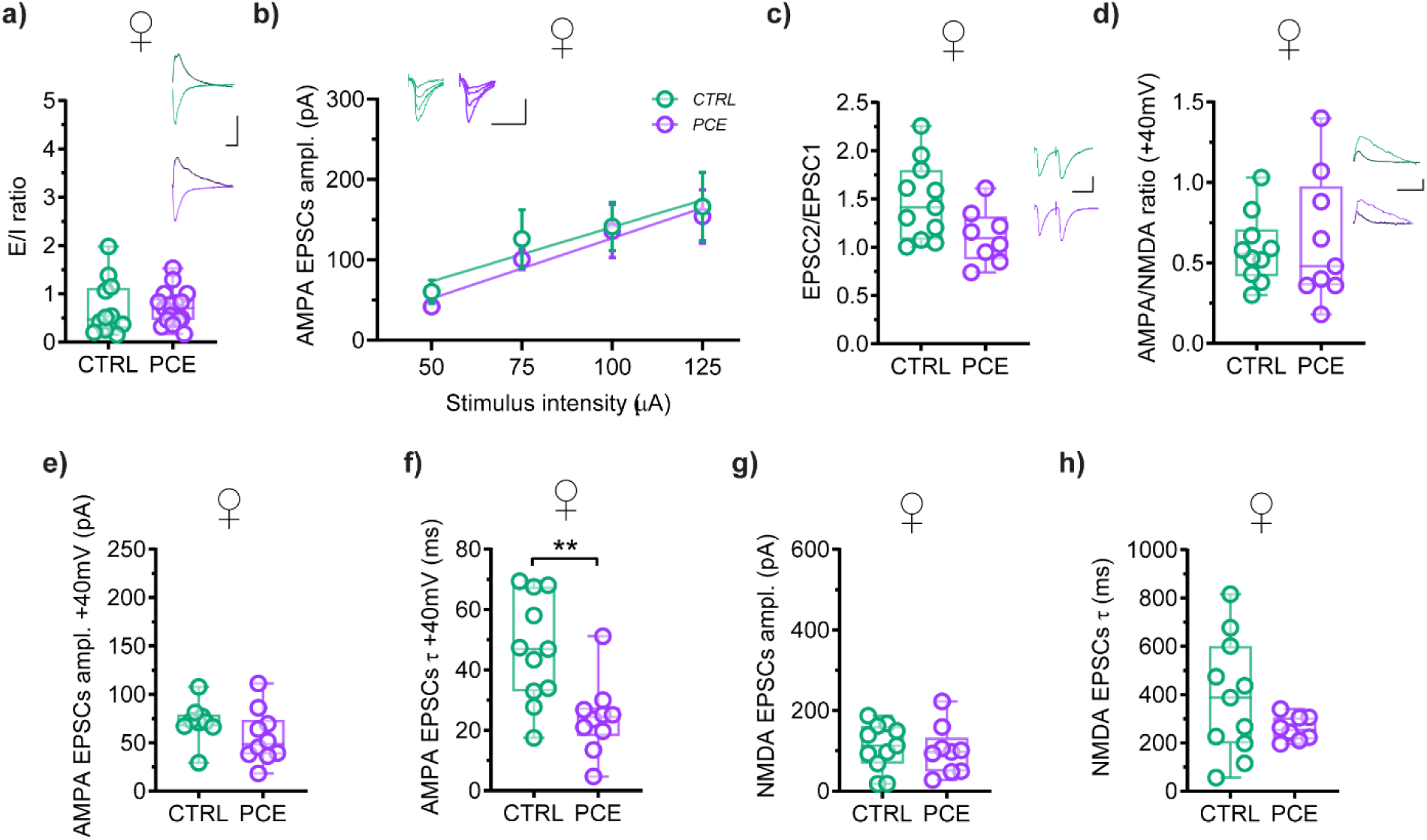
Effect of PCE on excitatory synaptic properties of PFC pyramidal neurons in preadolescent female offspring. (**a**) Plots and traces of evoked EPSCs (at -80 mV) and IPSCs (at 0 mV) recorded from PFC pyramidal neurons of CTRL and PCE female animals (p=0.4915; Mann–Whitney test; n_cells_=12-17/group, n_rats_=5-7/group); calibration bar: 25 ms, 200 pA. (**b**) Input-output relationships of AMPA-mediated EPSCs in females (interaction current x treatment p=0.9826; 2-way RM ANOVA; n_cells_=11-13/group; n_rats_=6-8/group). (**c**) Plots and traces of averaged paired-pulse ratio (EPSC2/EPSC1) of AMPA EPSCs of female pyramidal neurons at the higher stimulus intensity (n_cells_=11-13/group; n_rats_=6-8/group). Each symbol represents the averaged value (±S.E.M.) obtained from different cells. Insets show representative traces of AMPA EPSCs recorded from pyramidal neurons at each stimulus intensity; (**b,c**) calibration bar: 50 ms, 100 pA. (**d**) AMPA/NMDA ratio did not change in females (p=0.8868; Mann–Whitney test; n_cells_=9-10/group; n_rats_=6-7/group). Insets show representative traces of AMPA, and NMDA EPSCs recorded from pyramidal neurons held at +40 mV in PFC slices; calibration bar: 25 ms, 200 pA. (**e,f**) Quantification of the data showing AMPA EPSC amplitude (**e**; p=0.1333; Mann–Whitney test) and decay time kinetics (weighted tau, τ) (**f**; **p=0.0037; Mann–Whitney test) in CTRL and PCE female rats (n_cells_=9-11/group; n_rats_=6-7/group). (**g,h**) Quantification of the data showing NMDA EPSC amplitude (**g**) and τ (**h**) in CTRL and PCE female offspring (n_cells_=9-11/group; n_rats_=6-7/group).

When we examined the properties of GABA synaptic inputs on male mPFC pyramidal neurons, we found no effect on GABA_A_ IPSC amplitude (p=0.4104, 2-way RM ANOVA, interaction: F_(3,48)_=0.9792; **Fig. 6a**) and in the paired pulse ratio (U=19, p=0.1949, M-W test; **Fig. 6b**). Similarly, in females, PCE did not affect GABA synaptic efficacy (IPSCs amplitude: p=0.8354, 2-way RM ANOVA, interaction: F_(3,53)_=0.2858; paired-pulse ratio: U=27, p=0.1564, M-W test; **Fig. 6c-d**).

**Figure 6.**
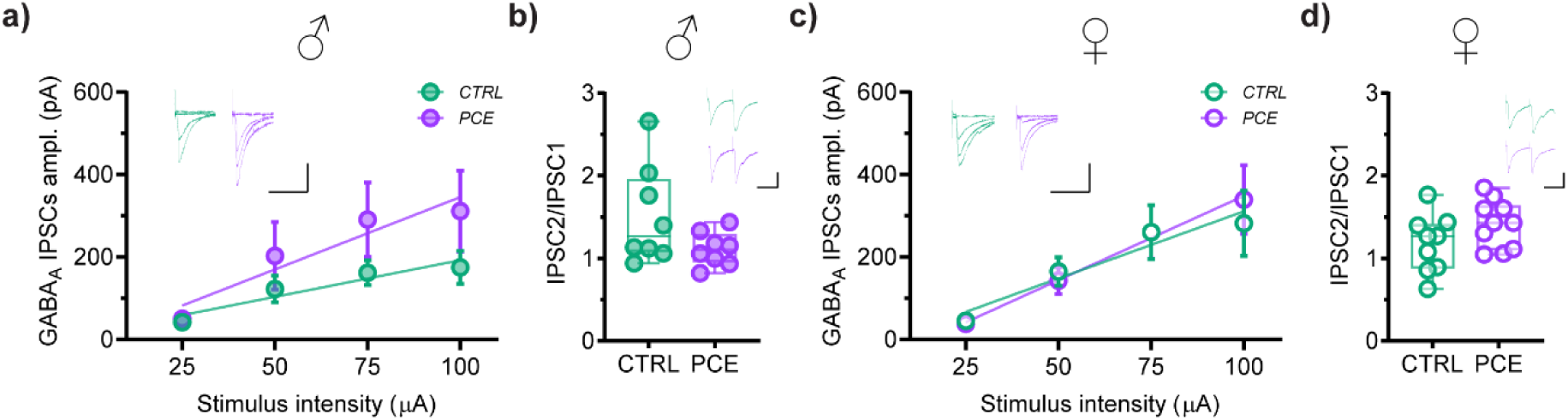
PCE did not alter synaptic inhibition onto pyramidal neurons in preadolescence. (**a**) Input-output relationships of GABA_A_-mediated IPSCs in males (interaction current x treatment p=0.4104; 2-way RM ANOVA; n_cells_=9-10/group; n_rats_=6-7/group). Each symbol represents the averaged value (±S.E.M.) obtained from different cells. (**b**) Plots of averaged paired-pulse ratio (IPSC2/IPSC1) of GABA_A_ IPSCs of pyramidal neurons at the higher stimulus intensity in male offspring (n_cells_=8/group; n_rats_=6-7/group). (**c**) Input-output relationships of GABA_A_-mediated IPSCs in females (interaction current x treatment p=0.8354; 2-way RM ANOVA; n_cells_=10/group; n_rats_=6/group). Each symbol represents the averaged value (±S.E.M.) obtained from different cells. (**d**) Plots of averaged paired-pulse ratio (IPSC2/IPSC1) of GABA_A_ IPSCs of pyramidal neurons at the highest stimulus intensity in female progeny (n_cells_=9-10/group; n_rats_=6/group). Calibration bar: 50 ms, 100 pA.

### 3.5 PCE effects on gene expression of endocannabinoid and dopaminergic systems in the prefrontal cortex of preadolescent rats

We next assessed whether PCE induced early molecular alterations in the PFC by examining the expression of genes involved in endocannabinoid and dopaminergic signaling in male preadolescent offspring. Our analysis revealed increased *Magl* mRNA levels in the PFC of male preadolescent PCE rats compared to controls (U=2, p=0.0087, M-W test, **Table 3**) with no changes observed in other components of endocannabinoid system (ECS; **Table 3**). Furthermore, *Dat* mRNA levels were also enhanced in the PFC of male preadolescent PCE rats compared to controls (U=0, p=0.0159, M-W test, **Table 3**) with no changes detected in dopaminergic receptor mRNA levels (*Drd1*, *Drd2* and *Drd3* encoding DA D1, D2 and D3 receptor, respectively, **Table 3**).

**Table 3.**
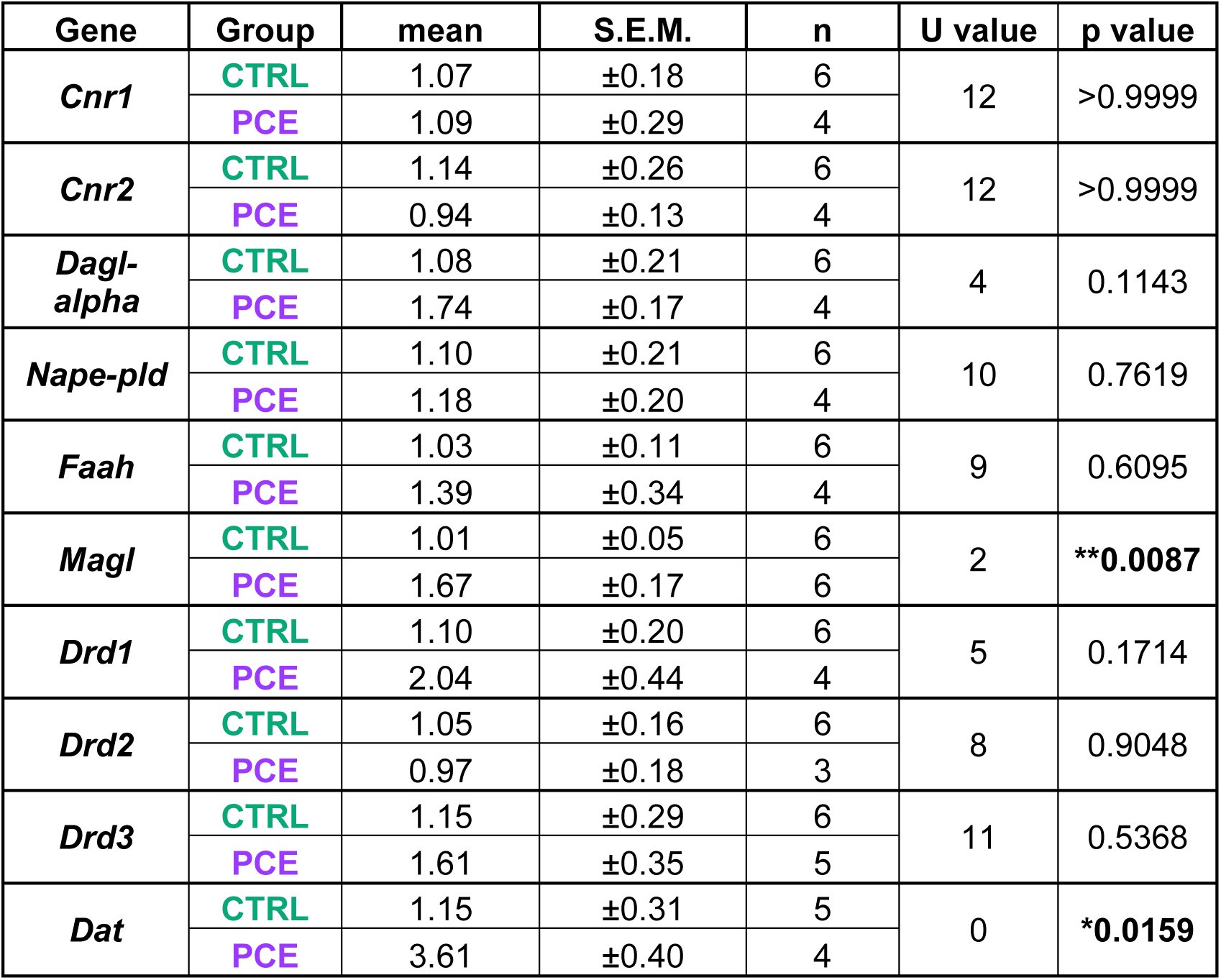
PCE increased Magl and Dat gene expression in the prefrontal cortex of male preadolescent rats. Gene expression data are reported as 2^-ΔΔCt^ values calculated by delta–delta Ct (ΔΔCt) method versus control (CTRL) rats. Expression was normalized to the mean of GAPDH and 18S housekeeping genes. Data are reported as mean ± S.E.M. p values, number (n) of rats per group and Mann-Whitney U values are indicated.

| Gene | Group | mean | S.E.M. | n | U value | p value |
| --- | --- | --- | --- | --- | --- | --- |
| *Cnr1* | CTRL | 1.07 | ±0.18 | 6 | 12 | >0.9999 |
|  | PCE | 1.09 | ±0.29 | 4 |  |  |
| *Cnr2* | CTRL | 1.14 | ±0.26 | 6 | 12 | >0.9999 |
|  | PCE | 0.94 | ±0.13 | 4 |  |  |
| *Dagl-alpha* | CTRL | 1.08 | ±0.21 | 6 | 4 | 0.1143 |
|  | PCE | 1.74 | ±0.17 | 4 |  |  |
| *Nape-pld* | CTRL | 1.10 | ±0.21 | 6 | 10 | 0.7619 |
|  | PCE | 1.18 | ±0.20 | 4 |  |  |
| *Faah* | CTRL | 1.03 | ±0.11 | 6 | 9 | 0.6095 |
|  | PCE | 1.39 | ±0.34 | 4 |  |  |
| *Magl* | CTRL | 1.01 | ±0.05 | 6 | 2 | **\*\*0.0087** |
|  | PCE | 1.67 | ±0.17 | 6 |  |  |
| *Drd1* | CTRL | 1.10 | ±0.20 | 6 | 5 | 0.1714 |
|  | PCE | 2.04 | ±0.44 | 4 |  |  |
| *Drd2* | CTRL | 1.05 | ±0.16 | 6 | 8 | 0.9048 |
|  | PCE | 0.97 | ±0.18 | 3 |  |  |
| *Drd3* | CTRL | 1.15 | ±0.29 | 6 | 11 | 0.5368 |
|  | PCE | 1.61 | ±0.35 | 5 |  |  |
| *Dat* | CTRL | 1.15 | ±0.31 | 5 | 0 | **\*0.0159** |
|  | PCE | 3.61 | ±0.40 | 4 |  |  |

Our analysis of female PFC also revealed increased *Magl* mRNA levels in PCE rats compared to controls (U=3.500, p=0.0173, M-W test, **Table 4**) with no changes detected in the expression of other genes.

**Table 4.** PCE increased Magl gene expression in the prefrontal cortex of female preadolescent rats. Gene expression data are reported as 2^-ΔΔCt^ values calculated by delta–delta Ct (ΔΔCt) method versus CTRL rats. Expression was normalized to the mean of GAPDH and 18S housekeeping genes. Data are reported as mean ± S.E.M. p values, n of rats per group and Mann-Whitney U values are indicated.

| Gene | Group | mean | S.E.M. | n | U value | p value |
| --- | --- | --- | --- | --- | --- | --- |
| *Cnr1* | CTRL | 1.10 | $\pm 0.23$ | 5 | 12.50 | 0.7056 |
| | PCE | 0.92 | $\pm 0.14$ | 6 | | |
| *Cnr2* | CTRL | 1.09 | $\pm 0.26$ | 5 | 12 | 0.6623 |
| | PCE | 1.07 | $\pm 0.22$ | 6 | | |
| *Dagl-alpha* | CTRL | 1.08 | $\pm 0.20$ | 6 | 15 | 0.6991 |
| | PCE | 1.12 | $\pm 0.06$ | 6 | | |
| *Nape-pld* | CTRL | 1.06 | $\pm 0.19$ | 5 | 14.50 | 0.9719 |
| | PCE | 0.98 | $\pm 0.11$ | 6 | | |
| *Faah* | CTRL | 1.12 | $\pm 0.21$ | 6 | 14.50 | 0.6147 |
| | PCE | 1.37 | $\pm 0.14$ | 6 | | |
| *Magl* | CTRL | 1.04 | $\pm 0.12$ | 6 | 3.500 | **\*0.0173** |
| | PCE | 1.67 | $\pm 0.17$ | 6 | | |
| *Drd1* | CTRL | 1.26 | $\pm 0.32$ | 6 | 6 | 0.1255 |
| | PCE | 2.10 | $\pm 0.39$ | 5 | | |
| *Drd2* | CTRL | 1.04 | $\pm 0.13$ | 6 | 11.50 | 0.3312 |
| | PCE | 0.85 | $\pm 0.14$ | 6 | | |
| *Drd3* | CTRL | 1.04 | $\pm 0.13$ | 5 | 7 | 0.1775 |
| | PCE | 1.57 | $\pm 0.25$ | 6 | | |
| *Dat* | CTRL | 1.08 | $\pm 0.18$ | 5 | 6 | 0.2222 |
| | PCE | 1.84 | $\pm 0.41$ | 5 | | |

### 3.6 PCE effect on Magl and Dat DNA methylation levels in the prefrontal cortex of preadolescent rats

To assess whether epigenetic mechanisms might account for the alterations observed in gene expression, we examined DNA methylation patterns within the regulatory regions of the *Magl* and *Dat* genes in the PFC of male and female preadolescent PCE and control rats. The schematic representation of rat *Magl* and *Dat* sequences analyzed is shown in **Figure 7**.

**Figure 7.**
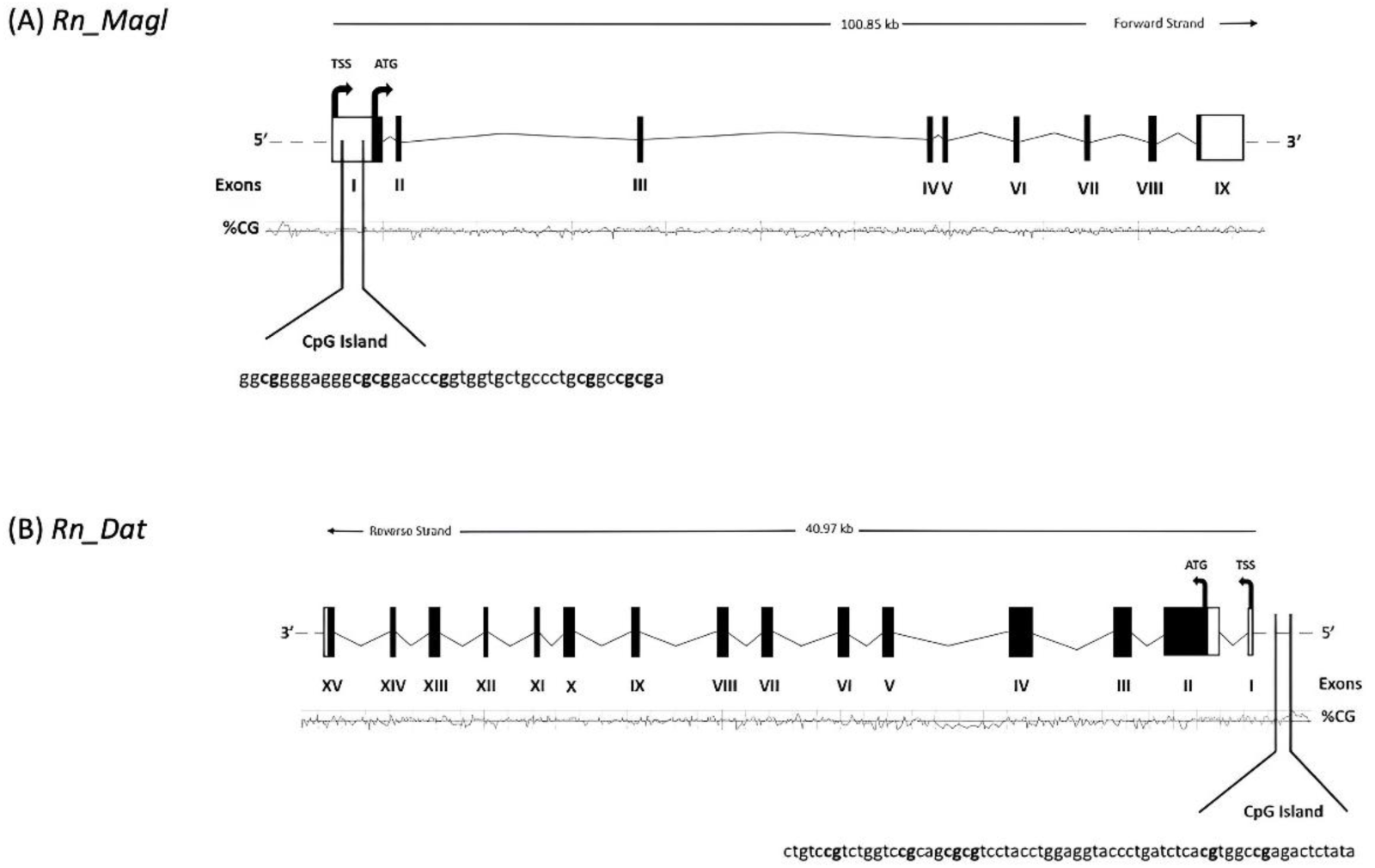
Schematic representation of rat (A) Magl (Transcript ID ENSRNOT00000110964.2, genome assembly Norway rat - BN/NHsdMcwi (GRCr8)) and (B) Dat (Transcript ID ENSRNOT00000040291.3, genome assembly Norway rat - BN/NHsdMcwi (GRCr8)) genes. ATG indicates the translation start site. Shown are the locations of CpG islands, exons, introns and the TSS (transcription start site). Coding regions of exons are shown darker shading. Analyzed sequences are indicated.

Results are detailed in Tables 5, 6 and 7,8 respectively.

**Table 5.** PCE did not affect Magl DNA methylation levels in the prefrontal cortex of male preadolescent rats. Comparison of DNA methylation status at the Magl gene regulatory region in the PFC of male preadolescent PCE and CTRL rats. DNA methylation data are presented as mean ± SEM of the methylation percentage values of individual CpG sites as well as of the average of the seven CpG sites under study. p values, n of rats per group and Mann-Whitney U values are indicated.

| *Magl* | Group | mean | S.E.M. | n | U value | p value |
| --- | --- | --- | --- | --- | --- | --- |
| CpG1 | CTRL | 0.88 | ±0.18 | 6 | 12 | >0.999999 |
|  | PCE | 0.90 | ±0.30 | 6 |  |  |
| CpG2 | CTRL | 1.44 | ±0.35 | 6 | 10.50 | 0.961491 |
|  | PCE | 1.85 | ±0.16 | 5 |  |  |
| CpG3 | CTRL | 2.20 | ±0.34 | 6 | 11 | 0.899262 |
|  | PCE | 1.71 | ±0.21 | 6 |  |  |
| CpG4 | CTRL | 3.45 | ±0.44 | 6 | 16.50 | 0.999782 |
|  | PCE | 3.68 | ±0.24 | 6 |  |  |
| CpG5 | CTRL | 1.71 | ±0.13 | 6 | 15.50 | 0.990228 |
|  | PCE | 1.97 | ±0.28 | 6 |  |  |
| CpG6 | CTRL | 3.37 | ±0.29 | 6 | 17.50 | >0.999999 |
|  | PCE | 3.49 | ±0.58 | 6 |  |  |
| CpG7 | CTRL | 2.35 | ±0.48 | 6 | 14 | 0.998589 |
|  | PCE | 2.00 | ±0.22 | 5 |  |  |
| Average | CTRL | 2.20 | ±0.11 | 6 | 12 | 0.999510 |
|  | PCE | 2.32 | ±0.21 | 5 |  |  |

**Table 6.** PCE did not affect Magl DNA methylation levels in the prefrontal cortex of female preadolescent rats. Comparison of DNA methylation status at the Magl gene regulatory region in the PFC of male preadolescent PCE and CTRL rats. DNA methylation data are presented as the mean ± SEM of the methylation percentage values of individual CpG sites as well as of the average of the seven CpG sites under study. p values, n of rats per group and Mann-Whitney U values are indicated.

| *Magl* | Group | mean | S.E.M. | n | U value | p value |
| --- | --- | --- | --- | --- | --- | --- |
| CpG1 | CTRL | 1.13 | ±0.25 | 6 | 17.50 | >0.999999 |
|  | PCE | 1.08 | ±0.26 | 6 |  |  |
| CpG2 | CTRL | 1.81 | ±0.20 | 6 | 13.00 | 0.942396 |
|  | PCE | 2.23 | ±0.33 | 6 |  |  |
| CpG3 | CTRL | 1.28 | ±0.35 | 6 | 15.00 | >0.999999 |
|  | PCE | 1.37 | ±0.46 | 6 |  |  |
| CpG4 | CTRL | 4.21 | ±0.61 | 6 | 13.00 | >0.999999 |
|  | PCE | 4.33 | ±0.62 | 5 |  |  |
| CpG5 | CTRL | 1.98 | ±0.29 | 6 | 13.00 | 0.985907 |
|  | PCE | 2.83 | ±0.37 | 6 |  |  |
| CpG6 | CTRL | 3.34 | ±0.82 | 6 | 13.00 | 0.999838 |
|  | PCE | 2.88 | ±0.50 | 5 |  |  |
| CpG7 | CTRL | 1.52 | ±0.34 | 6 | 7.000 | 0.595710 |
|  | PCE | 2.35 | ±0.32 | 6 |  |  |
| Average | CTRL | 2.18 | ±0.21 | 6 | 14.00 | >0.999999 |
|  | PCE | 2.24 | ±0.31 | 5 |  |  |

**Table 7.**
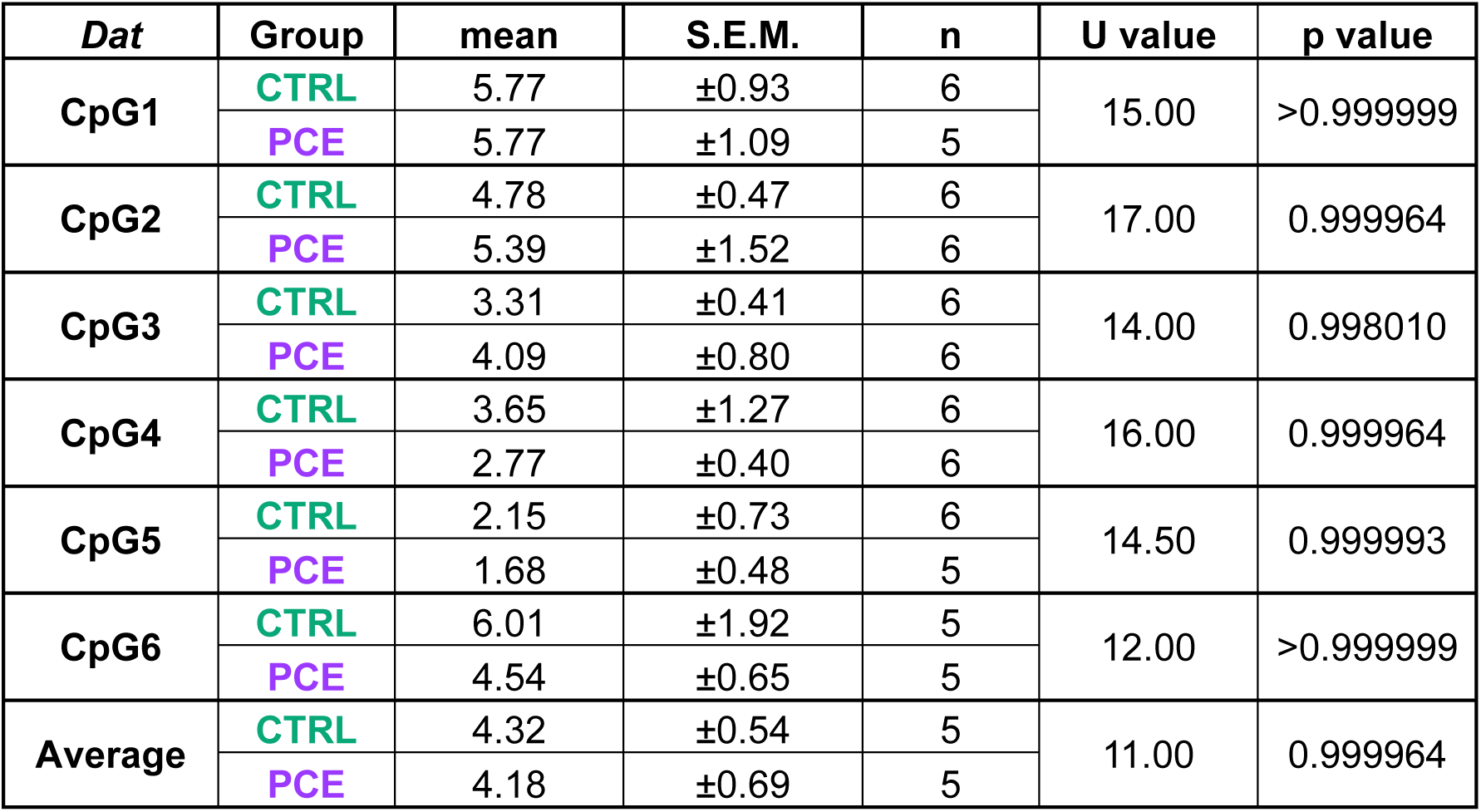
PCE did not affect Dat DNA methylation levels in the prefrontal cortex of male preadolescent rats. Comparison of DNA methylation status at the Dat gene regulatory region in the PFC of male preadolescent PCE and CTRL rats. DNA methylation data are presented as the mean ± SEM of the methylation percentage values of individual CpG sites as well as of the average of the six CpG sites under study. p values, n of rats per group and Mann-Whitney U values are indicated.

| *Dat* | Group | mean | S.E.M. | n | U value | p value |
| --- | --- | --- | --- | --- | --- | --- |
| CpG1 | CTRL | 5.77 | $\pm 0.93$ | 6 | 15.00 | >0.999999 |
| | PCE | 5.77 | $\pm 1.09$ | 5 | | |
| CpG2 | CTRL | 4.78 | $\pm 0.47$ | 6 | 17.00 | 0.999964 |
| | PCE | 5.39 | $\pm 1.52$ | 6 | | |
| CpG3 | CTRL | 3.31 | $\pm 0.41$ | 6 | 14.00 | 0.998010 |
| | PCE | 4.09 | $\pm 0.80$ | 6 | | |
| CpG4 | CTRL | 3.65 | $\pm 1.27$ | 6 | 16.00 | 0.999964 |
| | PCE | 2.77 | $\pm 0.40$ | 6 | | |
| CpG5 | CTRL | 2.15 | $\pm 0.73$ | 6 | 14.50 | 0.999993 |
| | PCE | 1.68 | $\pm 0.48$ | 5 | | |
| CpG6 | CTRL | 6.01 | $\pm 1.92$ | 5 | 12.00 | >0.999999 |
| | PCE | 4.54 | $\pm 0.65$ | 5 | | |
| Average | CTRL | 4.32 | $\pm 0.54$ | 5 | 11.00 | 0.999964 |
| | PCE | 4.18 | $\pm 0.69$ | 5 | | |

**Table 8.** PCE did not affect Dat DNA methylation levels in the prefrontal cortex of female preadolescent rats. Comparison of DNA methylation status at the Dat gene regulatory region in the PFC of male preadolescent PCE and CTRL rats. DNA methylation data are presented as the mean ± SEM of the methylation percentage values of individual CpG sites as well as of the average of the six CpG sites under study. p values, n of rats per group and Mann-Whitney U values are indicated.

| *Dat* | Group | mean | S.E.M. | n | U value | p value |
| --- | --- | --- | --- | --- | --- | --- |
| CpG1 | CTRL | 4.39 | $\pm 0.51$ | 6 | 14.50 | 0.978939 |
| | PCE | 4.46 | $\pm 0.96$ | 6 | | |
| CpG2 | CTRL | 3.14 | $\pm 0.86$ | 6 | 5.000 | 0.254698 |
| | PCE | 5.75 | $\pm 0.67$ | 6 | | |
| CpG3 | CTRL | 2.83 | $\pm 0.33$ | 6 | 17.50 | 0.978939 |
| | PCE | 2.32 | $\pm 0.80$ | 6 | | |
| CpG4 | CTRL | 4.76 | $\pm 1.67$ | 6 | 14.50 | 0.978939 |
| | PCE | 2.59 | $\pm 0.66$ | 6 | | |
| CpG5 | CTRL | 1.33 | $\pm 0.71$ | 6 | 9.500 | 0.718565 |
| | PCE | 3.16 | $\pm 1.28$ | 6 | | |
| CpG6 | CTRL | 4.70 | $\pm 0.73$ | 6 | 15.00 | 0.978939 |
| | PCE | 5.16 | $\pm 1.38$ | 6 | | |
| Average | CTRL | 3.53 | $\pm 0.49$ | 6 | 12.50 | 0.934315 |
| | PCE | 3.91 | $\pm 0.41$ | 6 | | |

DNA methylation analysis of each individual CpG site, as well as the average of all CpG sites analyzed within the *Magl* (**Tables 5-6**) and *Dat* (**Tables 7-8**) gene regulatory regions, revealed no significant differences between groups in the PFC.

### 3.7 PCE effects on TH and Netrin-1 immunoreactivity in the medial prefrontal cortex and nucleus accumbens of preadolescent rats

To assess whether PCE-induced PFC functional changes could reflect alterations in dopaminergic innervation, we measured the intensity of tyrosine hydroxylase (TH) immunoreactivity in the mPFC and in the nucleus accumbens (NAc; **Fig. 8a**). No differences in TH immunoreactivity were observed in the mPFC (2-way ANOVA, PCE effect F_(1, 27)_=0.00594; p=0.8092; sex effect F_(1, 27)_=1.02; p=0.3215; **Fig. 8b**), whereas this increased in the NAc as a function of PCE and irrespective of sex (2-way ANOVA, PCE effect F_(1, 26)_=8.319; p=0.0078; sex effect F_(1, 26)_=0.01894; p=0.8916; **Fig.8c**).

**Figure 8.**
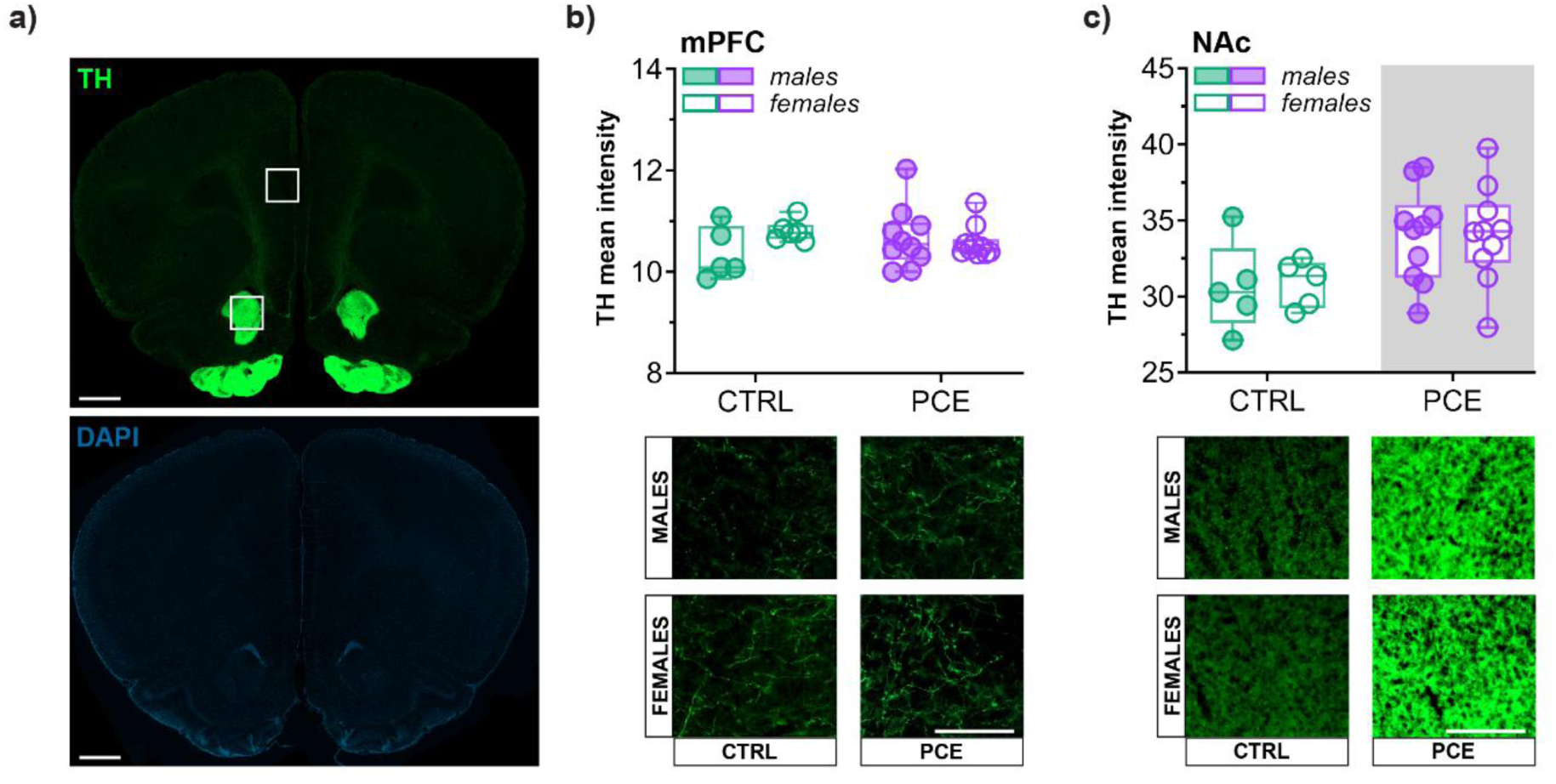
**Effect of PCE on TH immunoreactivity in the medial prefrontal cortex (mPFC) and nucleus accumbens (NAc)**. (**a**) Representative fluorescent images of tyrosine hydroxylase (TH) staining (green, top) and DAPI (blue, bottom) in a coronal brain slice, with the areas of interest for the mPFC and NAc. Scale bar, 1000 μm. (**b**) Graph and related images showing the density of TH immunoreactivity in the mPFC of CTRL and PCE male and female rats (no significant differences). Data are reported as mean ±SEM, n_rats_=5-10/group. Scale bar, 100 μm (**c**) Graph and related images showing the density of TH immunoreactivity in the NAc of CTRL and PCE male and female rats. (2-way ANOVA, PCE effect F_(1, 26)_=8.319; p=0.0078). Data are reported as mean ±SEM, n_rats_=6-10/group. Grey panel p<0.05. Scale bar, 100 μm. mPFC: medial prefrontal cortex; NAc: nucleus accumbens.

Given the role of axon guidance mechanisms in segregating dopaminergic projections during development, we next investigated the possibility that PCE could affect the Netrin-1/Dcc guidance cue system (Vosberg et al. 2020; Avramescu et al. 2024). We found that, in the PFC, Netrin-1 levels display sex differences, with female preadolescent rats showing higher levels than males, which were blunted by PCE (2-way ANOVA, sex effect F_(1, 28)_=4.901; p=0.0352; PCE effect F_(1, 28)_=2.905; p=0.0994; **Fig. 9a-c**). When we compared Netrin-1 expression levels in the NAc, we found no differences in the values as a function of sex or PCE (2-way ANOVA, sex effect F_(1, 28)_=1.862; p=0.1833; PCE effect F_(1, 28)_=0.7899; p=0.3817; **Fig. 9a, d, e**). Similarly, *Dcc* gene expression in the VTA displayed no effects of sex or PCE (2-way ANOVA, sex effect F_(1, 18)_=0.4076; p=0.5312; PCE effect F_(1, 18)_=1.344; p=0.2614; **Fig. 9f**).

**Figure 9.**
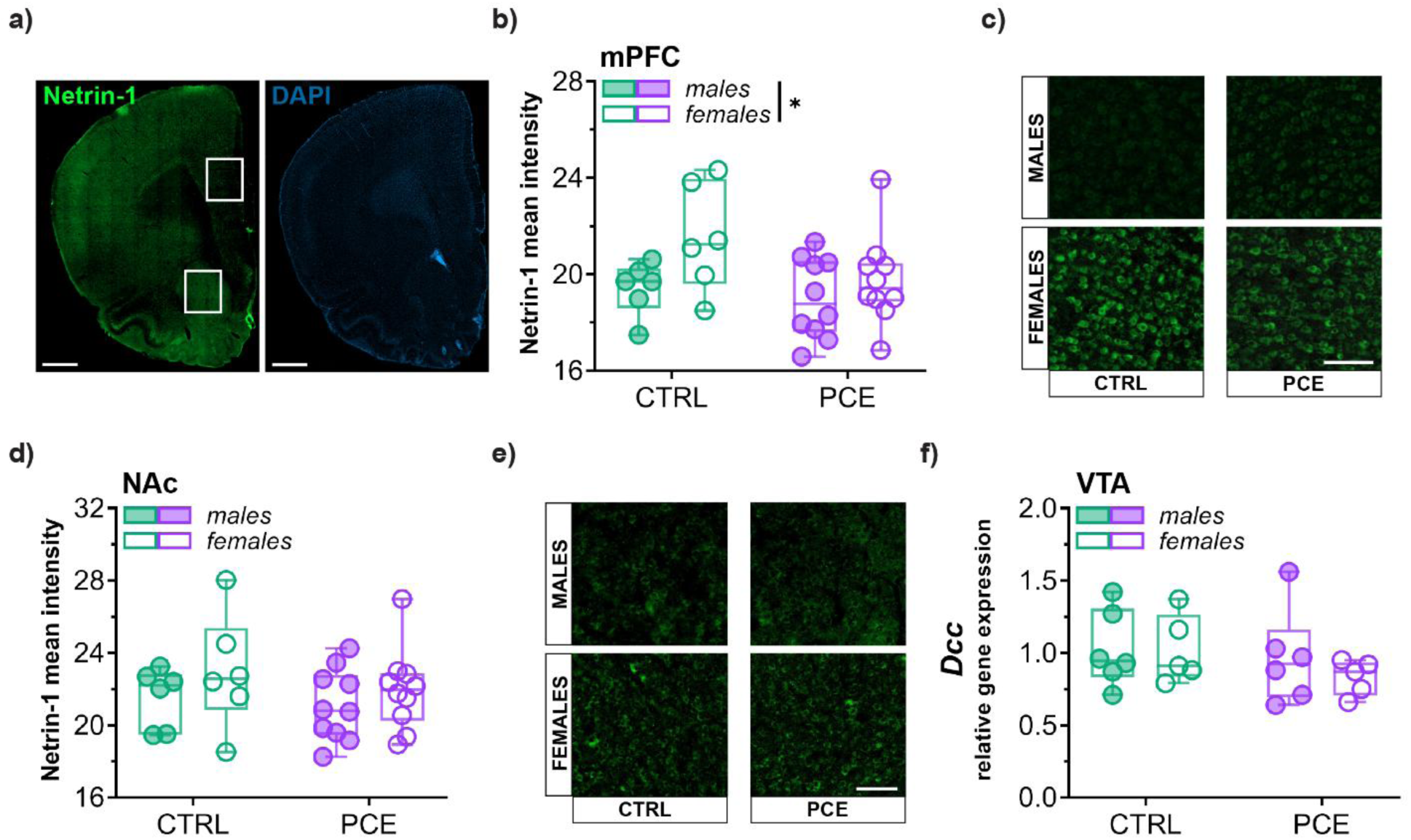
PCE effects on Netrin-1/Dcc signaling pathway. (**a**) Representative fluorescent images of Netrin-1 staining (green, left) and DAPI (blue, right) in a coronal brain hemi-slice, with the areas of interest for the mPFC and NAc. Scale bar, 1000 μm. (**b**) Graph and (**c**) related images showing the density of Netrin-1 immunoreactivity in the mPFC of CTRL and PCE male and female rats (2-way ANOVA, sex effect F_(1, 28)_=4.901; p=0.0352). Data are reported as mean ±SEM, n_rats_=6-10/group. Scale bar, 100 μm. (**d**) Graph and (**e**) related images showing the density of Netrin-1 immunoreactivity in the NAc of CTRL and PCE male and female rats. No significant differences were detected. Data are reported as mean ±SEM, n_rats_=6-10/group. Scale bar, 100 μm. (**f**) Dcc relative gene expression analyzed in the VTA of CTRL and PCE male and female preadolescent rats. No significant differences were detected. Data are reported as mean ±SEM, n_rats_=5-6/group. mPFC: medial prefrontal cortex; NAc: nucleus accumbens; VTA: ventral tegmental area.

## 4. Discussion

In the present study, we provide evidence that prenatal cannabinoid exposure (PCE) induces early, sex-specific alterations in behavioral, synaptic, and molecular features of the preadolescent prefrontal cortex (PFC). PCE increased marble-burying behavior, the excitability of PFC pyramidal neurons, and AMPA-mediated excitatory transmission selectively in male offspring. These changes occurred in the absence of detectable alterations in extracellular dopamine (DA) and 3,4-dihydroxyphenylacetic acid (DOPAC) mPFC extracellular levels or tyrosine hydroxylase (TH) immunoreactivity in the mPFC, suggesting that the behavioral phenotype is unlikely to reflect a gross alteration of tonic mesocortical DA availability. At the molecular level, PCE upregulated PFC *Magl* expression in both sexes while increasing *Dat* expression selectively in males, without detectable changes in DNA methylation at the analyzed regulatory regions. Finally, PCE blunted physiological sex differences in mPFC Netrin-1 immunoreactivity, pointing to an additional sex-specificity on developmental guidance-related signals.

Our findings corroborate the assumption that preadolescence is an early developmental window in which PCE is associated with male-specific behavioral alterations. Importantly, preadolescence precedes the full emergence of adolescent and adult behavioral phenotypes, while the PFC and its dopaminergic and endocannabinoid modulatory systems are still undergoing maturation (Schneider 2013; Caballero et al. 2016; Peters and Naneix 2022); therefore, this developmental window may be particularly informative for identifying early behavioral and neurobiological markers of vulnerability to later neuropsychiatric-like outcomes. Our evidence that PCE elicits repetitive behaviors during preadolescence supports an early onset of such behavior that manifests throughout adulthood (Mari et al. 2025), and that at this developmental stage may serve as a prodromic indicator for the onset of a variety of psychopathologies. Although marble burying behavior reflects species-typical digging, burying and digging can be sometimes dissociated (Thomas et al. 2009). While digging behavior *per se* might be interpreted as a natural behavior, marble burying is more representative of repetitive and perseverative behaviors (Dixit et al. 2020). Under our experimental conditions, this phenotype was observed specifically in male PCE offspring. This finding is consistent with previous evidence that developmental exposure to phytocannabinoids can alter marble-burying behavior, although both the direction and the sex specificity of this effect appear to depend on the experimental model. Indeed, perinatal exposure to Δ^9^-tetrahydrocannabinol (THC) or cannabidiol (CBD) increased marble burying selectively in adult female mice (Maciel et al. 2022), whereas a recent investigation reported a sex-neutral effect of prenatal exposure to either THC or CBD (Cáceres-Rodríguez et al. 2025). Thus, available findings suggest that prenatal/perinatal cannabinoid exposure may alter repetitive/perseverative behavioral domains in a context-dependent manner, with sex influences on the outcome depending on experimental conditions, such as species, developmental stages, time exposure windows, and type of cannabinoid administered.

Since the mesocortical DA system plays a pivotal role in the ability to suppress repetitive behaviors (Eagle et al. 2011; Wulaer et al. 2021; Fallon et al. 2013), one would expect a male-specific reduced mesocortical dopaminergic tone. However, our microdialysis observations argue against a gross alteration of mesocortical DA availability as a mechanism underlying the increased marble burying. The lack of detectable changes in extracellular DA levels in the PFC does not preclude its involvement in the PCE phenotype. Indeed, while the sampling rates required for *in vivo* microdialysis are optimized for detecting tonic neurotransmitter levels, they are often insufficient to capture the rapid, sub-second phasic DA transients that might drive the initiation and execution of a marble-burying task. Accordingly, we cannot rule out the possibility that the heightened *Dat* gene expression in PCE males might represent a homeostatic compensatory mechanism. Specifically, a PCE-induced increased dopaminergic tone may be effectively countered by elevated pools of DAT clearing a potential *surplus* of DA within the PFC, which would keep basal and THC-evoked tonic extracellular DA concentrations within physiological ranges. If so, enhanced DA reuptake might blunt the phasic DA signal needed to suppress repetitive behaviors (Popescu et al. 2016; Dreher and Burnod 2002), thereby contributing to the increased marble-burying phenotype observed in PCE males. Since DAT is crucial for regulating dopaminergic neurotransmission by controlling DA reuptake from synaptic clefts (Li et al. 2024), thereby playing a key role in behavioral regulation processes (Speranza et al. 2021), the observed upregulation of *Dat* expression may represent an adaptive mechanism engaged in response to altered dopaminergic signaling during development, which may potentially contribute to repetitive behavior. These changes are also consistent with our previous findings on transcriptional regulation of *Drd1* and *Drd2* in the PFC of PCE adolescent offspring (Di Bartolomeo et al. 2025) as well as of the *Drd2* in the PFC of adult male rats perinatally exposed to THC (Di Bartolomeo et al. 2021, 2023), and in peripheral blood of subjects with schizophrenia (Di Bartolomeo et al. 2021). Human studies have shown that PCE causes an upregulation of *Magl* alongside a decrease in *Dagl* gene expression in the fetal brain, ultimately leading to diminished endocannabinoid availability (Tortoriello et al. 2014). Not only our current observations that PCE upregulates *Magl* are in line with this finding, but are also consistent with our previous investigations in which we reported changes in endocannabinoid system (ECS) transcriptional regulation, specifically in the *Cnr1* gene in the PFC of adult rats perinatally exposed to THC (Di Bartolomeo et al. 2021) as well as in those exposed during both perinatal and adolescent phases (Di Bartolomeo et al. 2023) and in peripheral blood of subjects affected by schizophrenia (D’Addario et al. 2017). *Magl* is the main enzyme responsible for 2-arachidonoylglycerol (2-AG) degradation (Blankman et al. 2007); therefore, increased *Magl* expression may point to an early PCE-induced shift in 2-AG metabolism within the developing PFC (Wu et al. 2011; Calvigioni et al. 2014; Worley et al. 2020). The lack of methylation changes despite the upregulation of *Magl* and *Dat* gene expression suggests that other epigenetic mechanisms (e.g., histone modifications, chromatin remodeling, or non-coding RNA regulation) may account for these PCE-dependent effects. Accordingly, prenatal THC exposure can affect histone methyltransferases such as *Kmt2a* in the NAc of adult male rats (Ellis et al. 2022) as well as miRNAs expression in adult rat ovary (Martínez-Peña et al. 2022). Within this neurodevelopmental framework, our findings are compatible with the view that PCE may act as an early environmental insult capable of altering endocannabinoid-related maturation of prefrontal circuits. In agreement with the “two-hit” hypothesis of schizophrenia, our data extend the knowledge on PCE being a prenatal insult that disrupts endocannabinoid homeostasis long before adulthood (Frau et al. 2019b; Frau and Melis 2023). This early-life dysregulation mirrors the ECS abnormalities documented in schizophrenic patients, likely lowering the threshold for subsequent environmental insults to trigger neuropsychiatric onsets later in life, and positioning PCE as one of the most relevant environmental risk factors priming PFC architecture for subsequent disorders. Alternatively, developmental timing (Pagani et al. 2026; Schwarz et al. 2026) might make the difference: for instance, methylation patterns may have already been established earlier in development and been normalized by the preadolescent stage; or, conversely, methylation patterns may be established later during adulthood and not be observable at this stage of development. Further epigenetic mechanisms may be, therefore, investigated to clarify the transcriptional regulation of *Magl* and *Dat* genes driven by PCE. Consistent with the lack of detectable changes in extracellular DA and DOPAC levels, TH density was not changed in the mPFC of PCE progeny, whereas it increased in the *nucleus accumbens* (NAc) as a function of PCE. These differences in TH density between target regions persist throughout adolescence in PCE offspring (Di Bartolomeo et al. 2025) where a hyper-responsiveness to reward related cues, such as food- and drug- predictive cues, is detected (Luján et al. 2024). Future studies are warranted to directly assess DAT function and phasic DA dynamics and to clarify the contribution of dopaminergic mechanisms to this phenotype.

Our electrophysiological data reveal that the enhanced PFC excitability reported at adolescence (Di Bartolomeo et al. 2025) and adulthood (Bara et al. 2018) in male, but not female, rats exposed *in utero* to cannabinoids is already manifest at preadolescence. Such a hyper-excitable PFC may contribute to the failure in providing the inhibitory control required to suppress repetitive actions and to modulate emotional output, potentially leading to the emergence of compulsive-like behaviors observed here in male offspring. Such enhanced male PFC pyramidal excitability along with a larger maximal AMPA excitatory post synaptic current (EPSC) amplitude, are reminiscent of the electrophysiological profile of cortical neurons derived from ASD patients (Hussein et al. 2023). Our observation of a male-specific potentiation of AMPA signaling onto PFC pyramidal neurons is suggestive of synaptic potentiation (West and Atwood 2026; Wu et al. 2018; de León-López et al. 2025). PCE-potentiated AMPA/NMDA ratio might also reflect exposure-related neuroadaptations, analogous to the effects observed in the ventral tegmental area (VTA) DA neurons of preadolescent offspring exposed *in utero* to THC (Frau et al. 2019b). The observation of a sex-modulated PCE effect on AMPA EPSC decay time in PFC pyramidal neurons suggests that PCE-induced adaptations may follow distinct developmental trajectories depending on sex and age.

Our observation on the Netrin-1/DCC pathway, key in development and maturation of DA circuits (Vosberg et al. 2020; Reynolds et al. 2018; Cuesta et al. 2020), support and extend the studies showing that altered Netrin-1/DCC signaling can be affected by early life drug exposure (Reynolds et al. 2015, 2023; Hoops and Flores 2017). Hence, this may represent an additional developmental mechanism through which PCE affects prefrontal and mesolimbic maturation. While our data show that PCE did not modify *Dcc* mRNA levels in the VTA or Netrin-1 immunoreactivity in the NAc at preadolescence, we previously found changes at late adolescence (Di Bartolomeo et al. 2025), thus supporting the importance of longitudinal studies. Notably, and for the first time to our knowledge, we here describe that mPFC Netrin-1 immunoreactivity exhibits a sex dimorphism, with females displaying higher levels than males. This female-specific pattern is consistent with our previous findings in the mPFC of late adolescent rats (Di Bartolomeo et al. 2025). In contrast, not only mice do not show sex differences in Netrin-1 levels, but these were also higher in the PFC when compared to NAc. Hence, our findings suggest that Netrin-1 expression may be regulated in a species-, region-, and developmental-stage- specific manner (Reynolds et al. 2023). Because behavioral and electrophysiological phenotypes were male-specific, the functional relevance of the mPFC Netrin-1 pattern remains to be clarified.

Collectively, our findings reveal that PCE induces male-specific neurobehavioral dysfunction in the PFC of preadolescent rats. Enhanced repetitive behavior is accompanied by a heightened neuronal excitability of PFC pyramidal neurons, and an upregulation of two genes involved dopaminergic and ECS signaling pathways. Finally, these data suggest that PCE-induced behavioral impairments may be associated with complex changes involving both ECS and dopaminergic homeostasis in the developing male PFC.

## Author contributions

Conceptualization, M.M. and C.D.; methodology, S.A., M.D.B., V.S., F.T., P.D., M.C. and R.F.; formal analysis, S.A., M.D.B., F.T., M.C., G.L., P.D. and M.S.; investigation, S.A., M.D.B., V.S., F.T., M.C., G.L., M.S., P.D., P.S. and M.P.; data curation, S.A., M.D.B., M.C., M.M. and C.D.; writing—original draft preparation, S.A., M.D.B., M.C., M.M. and C.D.; writing—review and editing, S.A., M.D.B., M.C., V.S., M.S., G.L., M.P., R.F., M.M. and C.D.; supervision, M.M. and C.D.; funding acquisition, M.M. and C.D. All authors have read and agreed to the published version of the manuscript.

## Author approvals

This manuscript has not been accepted or published elsewhere. All authors have read and approved the final manuscript.

## Ethical statements

All procedures were performed in accordance with the European legislation (EU Directive, 2010/63/EU) and were approved by the Animal Ethics Committees of the University of Cagliari and by Italian Ministry of Health (auth. n. 636/2022-PR). The manuscript does not contain clinical studies or patient data.

## Data availability statement

Data can be provided from the corresponding authors upon reasonable request.

